# Annotation of snoRNA abundance across human tissues reveals complex snoRNA-host gene relationships

**DOI:** 10.1101/2021.02.11.430834

**Authors:** Étienne Fafard-Couture, Danny Bergeron, Sonia Couture, Sherif Abou Elela, Michelle S Scott

## Abstract

**Background:** Small nucleolar RNAs (snoRNAs) are mid-size non-coding RNAs required for ribosomal RNA modification, implying a ubiquitous tissue distribution linked to ribosome synthesis. However, increasing numbers of studies identify extra-ribosomal roles of snoRNAs in modulating gene expression, suggesting more complex snoRNA abundance patterns. Therefore, there is a great need for mapping the snoRNome in different human tissues as the blueprint for snoRNA functions.

**Results:** We used a low structure bias RNA-Seq approach to accurately quantify snoRNAs and compare them to the entire transcriptome in seven healthy human tissues (breast, ovary, prostate, testis, skeletal muscle, liver and brain). We identified 475 expressed snoRNAs categorized in two abundance classes that differ significantly in their function, conservation level and correlation with their host gene: 390 snoRNAs are uniformly expressed and 85 are enriched in the brain or reproductive tissues. Most tissue-enriched snoRNAs are embedded in lncRNAs and display strong correlation of abundance with them, whereas uniformly expressed snoRNAs are mostly embedded in protein-coding host genes and are mainly non- or anticorrelated with them. 59% of the non-correlated or anticorrelated protein-coding host gene/snoRNA pairs feature dual-initiation promoters, as opposed to only 16% of the correlated non-coding host gene/snoRNA pairs.

**Conclusions:** Our results demonstrate that snoRNAs are not a single homogeneous group of housekeeping genes but include highly regulated tissue-enriched RNAs. Indeed, our work indicates that the architecture of snoRNA host genes varies to uncouple the host and snoRNA expressions in order to meet the different snoRNA abundance levels and functional needs of human tissues.

## Background

Small nucleolar RNAs (snoRNAs) are a conserved family of mid-size non-coding RNA best characterized as guides for the chemical modification of nascent ribosomal RNA (rRNA) leading to ribosome assembly [1–3]. Functional snoRNAs are a part of larger ribonucleoprotein complexes (snoRNPs) composed of core proteins required for snoRNA stability that represent an enzymatic moiety needed for the RNA modification reaction [2,4–6]. SnoRNAs are divided in two types based on their structure and the modification they catalyze. Box C/D snoRNAs interact with the methyltransferase fibrillarin and guide the 2’-O-methylation of their target RNA while box H/ACA snoRNAs bind the pseudouridine synthase dyskerin and catalyze pseudouridylation [4,7,8]. Recently, a small number of box C/D snoRNAs have been shown to guide the acetylation of rRNA [9,10]. In addition to rRNA, snoRNAs also guide modifications on small nuclear RNAs (snRNAs) and a small subset including SNORD3 (U3) and SNORD118 (U8) are involved in rRNA processing [1,11]. Other snoRNAs have no known target in rRNA or snRNAs and are referred to as “orphan” snoRNAs [8].

A growing number of orphan snoRNAs as well as snoRNAs with rRNA or snRNA targets are being assigned alternative functions in the regulation of gene expression including at the level of chromatin remodeling, pre-mRNA stability, alternative splicing and polyadenylation (reviewed in [3,8,12]). In most cases, snoRNAs regulate their targets through base-pairing with the target sequence. This pairing may occur either in *trans* as in the case of rRNA modification guides or through *cis* base pairing that modifies the local structure surrounding the snoRNA, a mechanism that may be involved in snoRNA biogenesis [13]. It can be noted that despite their name, not all snoRNAs function in the nucleolus, particularly those involved in non-canonical roles [14,15]. The importance of the regulatory roles of snoRNAs is becoming increasingly clear by their association with a multitude of human diseases (reviewed in [16–19]). The scope and breadth of snoRNA regulatory functions likely extends beyond the few currently documented examples, given the large number of orphan snoRNAs and the fact that some rRNA- and snRNA-guiding snoRNAs have also been shown to have gene expression regulatory functions [3,12]. This raises the question of how the expression of snoRNAs is controlled to support their non-canonical regulatory functions.

In human, with the exception of the few snoRNAs required for rRNA processing, the majority of snoRNAs are expressed from the introns of either protein-coding or non-coding host genes (HGs) (Figure 1A and [20]). Accordingly, the expression of most snoRNAs depends, at least theoretically, on the transcription and splicing of their HG [5,20]. However, recent studies have started to provide examples of snoRNAs that might be uncoupled from the expression of their HG and even one orphan snoRNA that could regulate the splicing of its HG as a function of the amount of protein produced by the host [13]. The main mechanism uncovered so far uses nonsense-mediated decay (NMD) that permits degradation of the host transcript while preserving the expression of the snoRNA [6,21]. The idea of uncoupled snoRNA/HG expression was recently supported by the study of acute myeloid leukemia (AML) cells, human ovarian cell lines and mouse cell types that displayed limited correlation between a snoRNA and its HG expression [22–24]. However, the uncoupling seen in these proliferative models could also be the result of a methylation and pseudouridylation system that is unable to keep up with increased ribosome biogenesis, leading to discrepancies between snoRNA and HG abundances. More recently, it was hypothesized that promoters with dual-initiation of transcription may provide means to separate the expression of snoRNA from that of the HGs [25]. Nonetheless, it is unclear if these heterogeneities in snoRNA and host expression are stochastic differences arising from variation in cell cultures or reflect a stable tissue-specific regulatory program.

**Figure 1.**
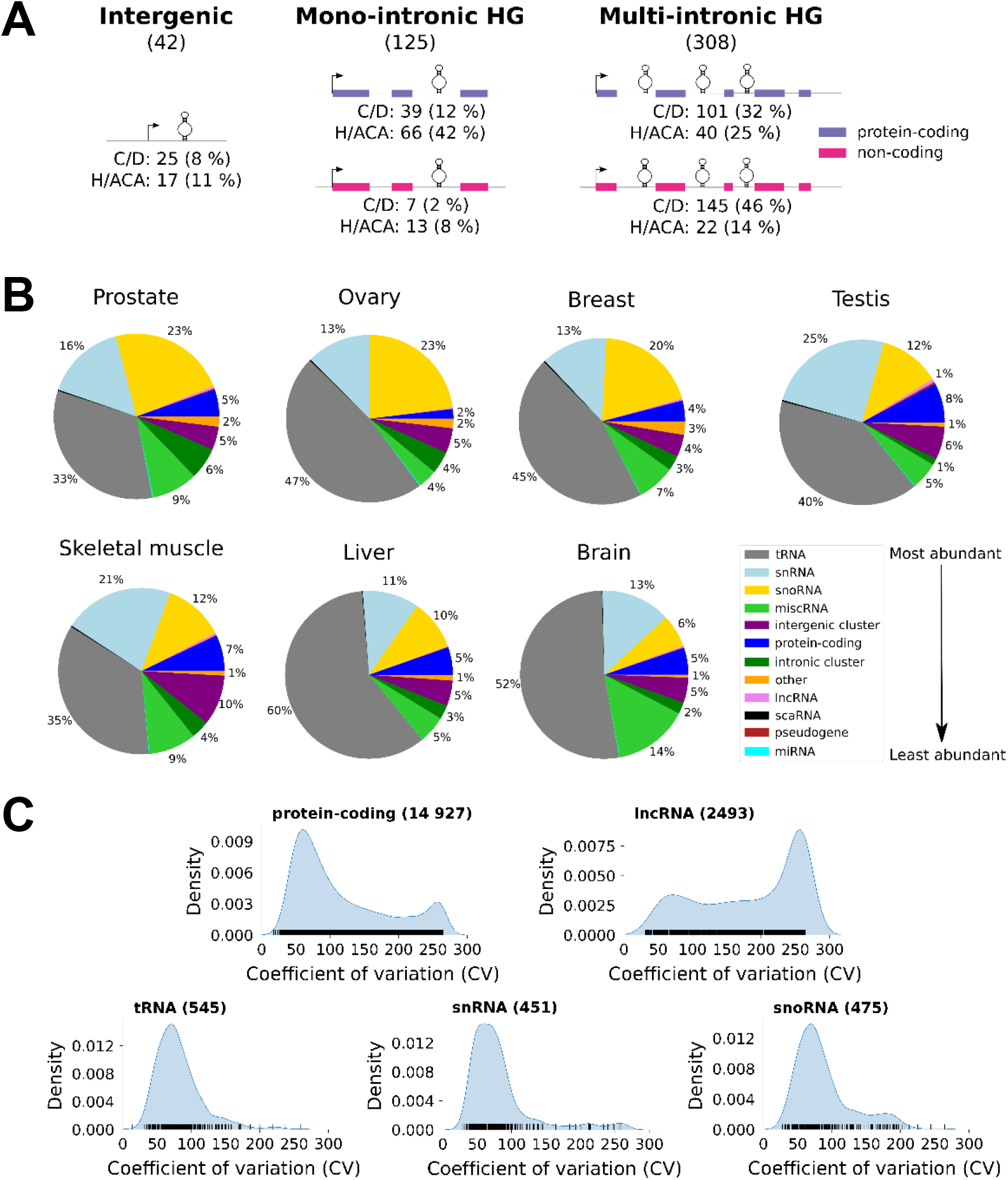
SnoRNAs are amongst the most abundant RNAs in the cell. (A) SnoRNAs are expressed from various genomic contexts. Shown is a schematic representation of the three most common classes of human snoRNA genomic contexts. Intergenic, mono-intronic HG and multi-intronic HG indicate respectively snoRNAs expressed as independent genes, host genes that encode a single snoRNA in one of their introns and host genes that encode multiple snoRNAs each in a separate intron. Protein-coding and non-coding host genes are indicated in purple and magenta respectively. The total number of expressed snoRNAs in each context is indicated under each title. The number and proportion of box C/D and H/ACA snoRNAs in each context are indicated under each schematic representation. The percentages are calculated for box C/D and H/ACA snoRNAs separately. (B) The highest relative abundance of snoRNAs is detected in prostate and female reproductive tissues. The RNA was sequenced using TGIRT-Seq from three replicates per tissue and the distribution of the average total abundance (in transcript per million (TPM)) per RNA biotype for each considered tissue is illustrated in the form of pie charts. Only RNAs with an abundance greater than 1 TPM in at least one tissue sample are considered. The color legend for the RNA biotype is shown on the far right, with an arrow representing the average ranking of RNA biotype abundance across tissues. (C) The abundance of snoRNAs is at the interface between regulatory RNAs and housekeeping RNAs. The distribution of snoRNA coefficient of variation (CV) was compared to that of the main classes of regulatory RNAs (protein-coding RNA and lncRNA) and housekeeping RNAs (tRNA and snRNA). The CV of each RNA is indicated by a vertical black line above the x-axis. The number of expressed RNAs considered in the density plots is indicated between parentheses on top of the graphs.

The most reported tissue-specific expression of snoRNA is found in the brain, where several snoRNAs were found to be predominantly expressed including the SNORD115 and SNORD116 families [26,27]. Despite these sporadic examples, the tissue distribution of the majority of the human snoRNome remains largely unexplored. Defining the human snoRNome is challenging due to the inherent difficulty in sequencing and quantifying the highly structured snoRNAs, especially when considered in relation to the abundance of their HG transcripts [23,28]. Indeed, the highly stable structure of snoRNAs impairs their reverse transcription, biasing most sequencing techniques towards the detection of less structured RNAs such as protein-coding transcripts [23,29]. The sequencing bias is not limited to non-snoRNA transcripts but is also detected between snoRNA types. Most sequencing techniques strongly favor the detection of box C/D snoRNAs over box H/ACA snoRNAs, presumably due to differences in the structure of these two snoRNA types (e.g. [22,27,30,31]). Aside from the reverse transcription sequencing bias, quantification errors are often encountered in assigning the snoRNA reads since the majority of snoRNAs exist in multiple copies and/or are embedded in introns, causing their reads to be either discarded or erroneously assigned to the HG [32,33].

Driven by the need to characterize the human snoRNome, we have used our newly developed snoRNA sensitive RNA-Seq pipeline [23] to measure the abundance of both snoRNAs and HG transcripts in seven healthy human tissues (breast, ovary, prostate, testis, skeletal muscle, liver and brain). By using a combination of thermostable group II intron reverse transcriptase sequencing (TGIRT-Seq) [23] and a read assignment pipeline that increases the accuracy of quantifying repeated and intron-embedded RNAs [33], we simultaneously followed the snoRNA and HG accumulations in the different tissues and provide a detailed portrait of the human snoRNome. Altogether, the results indicate that the abundance of snoRNAs is mostly defined by their genomic context and the architecture of their HG, which determines the level and type of tissue specificity and the degree of correlation between the snoRNA and HG abundance.

## Results

### Most expressed human snoRNAs are produced from intron-embedded genes

To determine the tissue distribution of snoRNAs and their relative abundance within the human transcriptome, we sequenced total ribodepleted fragmented RNA from seven healthy human tissues (breast, ovary, prostate, testis, skeletal muscle, liver and brain). Each tissue was sourced from 3 different individuals and sequenced using TGIRT-Seq methodology, which was shown to reliably quantify the abundance of different types of RNA in a same sample [23,29]. Indeed, in general our ranking of the abundance of RNAs was in agreement with the Genotype-Tissue Expression (GTEx) estimates for protein-coding genes (Tables S1A-G) [34]. The clustering of the quantified transcripts of all detected biotypes supports the quality of our datasets. Indeed, despite the expected differences between individuals and variations in sample composition, we notice little variability between samples of same tissue (Figure S1). Using this sequencing method, we detected RNA (>1 transcript per million (TPM) in at least one tissue sample) generated from 475 (50 %) snoRNA genes out of a total 947 annotated human snoRNA genes (Table S2). This is consistent with the fact that most RNAs are poorly expressed and only a minority of the transcriptome is highly expressed (Figure S1), as we have previously reported [23]. The majority (433 out of 475 snoRNAs, i.e. 91 %) of the expressed snoRNA genes are located in introns, while only 9 % (42 out of 475 snoRNAs) are located in intergenic regions and thus likely expressed from an independent promoter (Figure 1A). In contrast, 21 % of all annotated snoRNAs are located in intergenic regions, suggesting that most annotated intergenic snoRNA genes are not expressed. Indeed, intergenic snoRNAs contribute only to 2 % of the total snoRNA abundance, confirming the mostly intronic origin of human snoRNAs [35]. Interestingly, most expressed box H/ACA snoRNAs (67 %) are found in protein-coding HGs while expressed box C/D snoRNAs do not show clear HG biotype preference (Figure 1A). Variations in the number of snoRNA embedded in each HG is also observed between the two types of snoRNAs. The majority of box H/ACA snoRNAs (50 %) are the only snoRNA embedded within their HG (Figure 1A middle panel, mono-intronic HG), while the majority of box C/D snoRNAs (78 %) are encoded with multiple snoRNAs in separate introns of the same HG (Figure 1A right panel, multi-intronic HG). Together these results indicate that the two types of snoRNA have distinct embedding preferences.

### SnoRNAs are amongst the most abundant RNAs in the cell

To evaluate the relative contribution of snoRNAs to the transcriptome of the different human tissues, we compared their abundance to other RNA biotypes detected in each of the tissues examined. Overall, the highest percentage of expressed non-rRNA transcripts was detected within tRNAs where 84 % of the annotated genes are expressed at least in one tissue, followed by the protein-coding genes and snoRNA genes (Table S2). The lowest proportion of expressed genes was detected in the snRNA and lncRNA biotypes, which put the snoRNAs at the interface between translation associated RNAs and RNAs associated with RNA processing and regulation. Comparison of the number of transcripts (in TPM) generated from each biotype indicates that tRNA genes generate the highest number of transcripts regardless of the tissues examined (Figure 1B), which is in accordance with biochemical estimates [36]. On the other hand, the snoRNA and snRNA biotypes compete for the second place in the transcriptome in a tissue dependent manner. In the tissues derived from reproductive organs, except for testis, the snoRNAs are more abundant than snRNAs, while the snRNAs are more abundant in the other tissues, with the highest relative proportion of snRNA abundance detected in testis (Figure 1B). However, it is important to note that unlike snoRNAs, the snRNA transcripts are generated by only 24 % of the annotated snRNA genes and are driven by only a few genes that each generate more than 1000 TPMs like 7SK and spliceosomal snRNA genes (Table S2, Figures S2B and S3B). In contrast, half of the annotated snoRNAs generate around 15-20 % of non-rRNA transcripts which is half-way between the tRNAs at one extreme where 84 % of the annotated genes generated 45 % of transcripts and protein-coding RNAs where 73 % of the genes generate only 5 % of transcripts (Figure 1B and Table S2). In general, box C/D snoRNAs are on average 3 times more abundant than box H/ACA snoRNAs across tissues (Figure S4A). This ratio represents a lower abundance difference than what was previously reported between the two snoRNA types [22,27,30,31], which is likely explainable by the low structure bias approach we used. Nonetheless, both box C/D and H/ACA snoRNAs are mostly abundant to at least 1 TPM in all the studied tissues (Figure S4B), underlining the widespread importance of both snoRNA types in all human tissues. Overall, the abundance of most snoRNAs and tRNAs is more than 10 TPM in each tissue, whereas the abundance of other biotypes is mostly between 0 and 10 TPM (Figure S2). We conclude that on average snoRNA genes generate the highest diversity and number of non-rRNA transcripts after tRNAs in the human genome.

### Tissue-dependent distribution of RNA accumulation identifies two snoRNA abundance classes

In most cases, variations of RNA abundance are often taken as a basis for gene regulation and tissue specificity. Accordingly, we examined the pattern of snoRNA abundance in the different tissues and compared it to that of other RNA biotypes. As with snRNAs and tRNAs, the cumulative abundance curves seen with snoRNAs are less variable between tissues than those observed with protein-coding RNAs and lncRNAs (Figure S3), highlighting the widespread distribution of housekeeping RNAs across tissues. Of note, the most extreme examples of tissue specialization were observed in the case of the genes coding for albumin (ALB) and haptoglobin (HP), which produce as high as 20 % of all protein-coding transcripts in liver (Figure S3D). Similarly, most tissues express a very small number of lncRNAs except testis which is known for its permissive chromatin environment (Figure S3E) [37]. To enable direct comparison between the tissue distribution patterns of the different RNAs, we calculated the coefficient of variation (CV) for each RNA based on its abundance across the studied tissues (see Methods for more details). This metric allows us to numerically differentiate between the different degrees of tissue uniformity and enrichment of the different transcripts. Uniformly expressed RNAs are identified by low CV value, while tissue-enriched RNAs are identified by high CV value. Interestingly, comparison of the CV value of the different biotypes indicates that snoRNAs occupy a middle ground between the highly uniform tRNAs and snRNAs (CV<125) and highly variable protein-coding RNAs and lncRNAs (CV>125) (Figure 1C). In general, the uniformly expressed biotypes like tRNA and snRNA display a single peak with a median CV of around 65. In contrast, the tissue-enriched biotypes like protein-coding RNA and lncRNA display a bimodal distribution of CV, which indicates the presence of two RNA subpopulations, the first peak around a CV of 65 and the other around 260. Like the tissue-enriched protein-coding RNAs and lncRNAs, snoRNAs include two RNA subpopulations, the main one peaking at a CV of 70. However, unlike these tissue-enriched RNAs, the right-most snoRNA peak is much smaller and centered around a CV of 180. This bimodal distribution of snoRNA CVs can be split into two snoRNA abundance classes separated by a CV threshold of 125 (Figures 1C and S5; see Methods for more information). Accordingly, we termed the snoRNAs with a CV<125 “Uniformly expressed” or “UE” and snoRNAs with a CV>125 “Tissue-enriched” or “TE”. Taken together, these results indicate that snoRNA abundance is at the interface between that of housekeeping RNAs and regulatory RNAs, and that snoRNAs can be categorized into two distinct abundance classes.

### The majority of tissue-enriched snoRNAs are enriched in brain and reproductive tissues

To understand the origin and distribution of the two snoRNA abundance classes, we followed the accumulation of each RNA of these two classes in the different tissues. As indicated in Figure 2A, TE and UE snoRNAs generally clustered separately, validating the group identity of most RNA in each class. In addition, snoRNA abundance results in an adequate clustering of the tissues, once again confirming the validity of our datasets (Figure 2A). Analysis of individual snoRNA distribution indicates that the majority of snoRNAs (n=390) are uniformly expressed across tissues, whereas 85 snoRNAs are enriched in specific tissues (Figures 2B). Overall, 47 TE snoRNAs are enriched in the brain and 38 are enriched in male or female reproductive tissues (Figures 2A and S6B). The brain-enriched snoRNAs include the previously established brain-specific snoRNA family SNORD115 (Figure S6A) [26], validating our CV-based classification of snoRNAs. Interestingly, four snoRNAs with known rRNA targets (SNORA81, SNORA19, SNORD36A and SNORD111B) are highly enriched in both studied female reproductive tissues (Figure S6).

**Figure 2.**
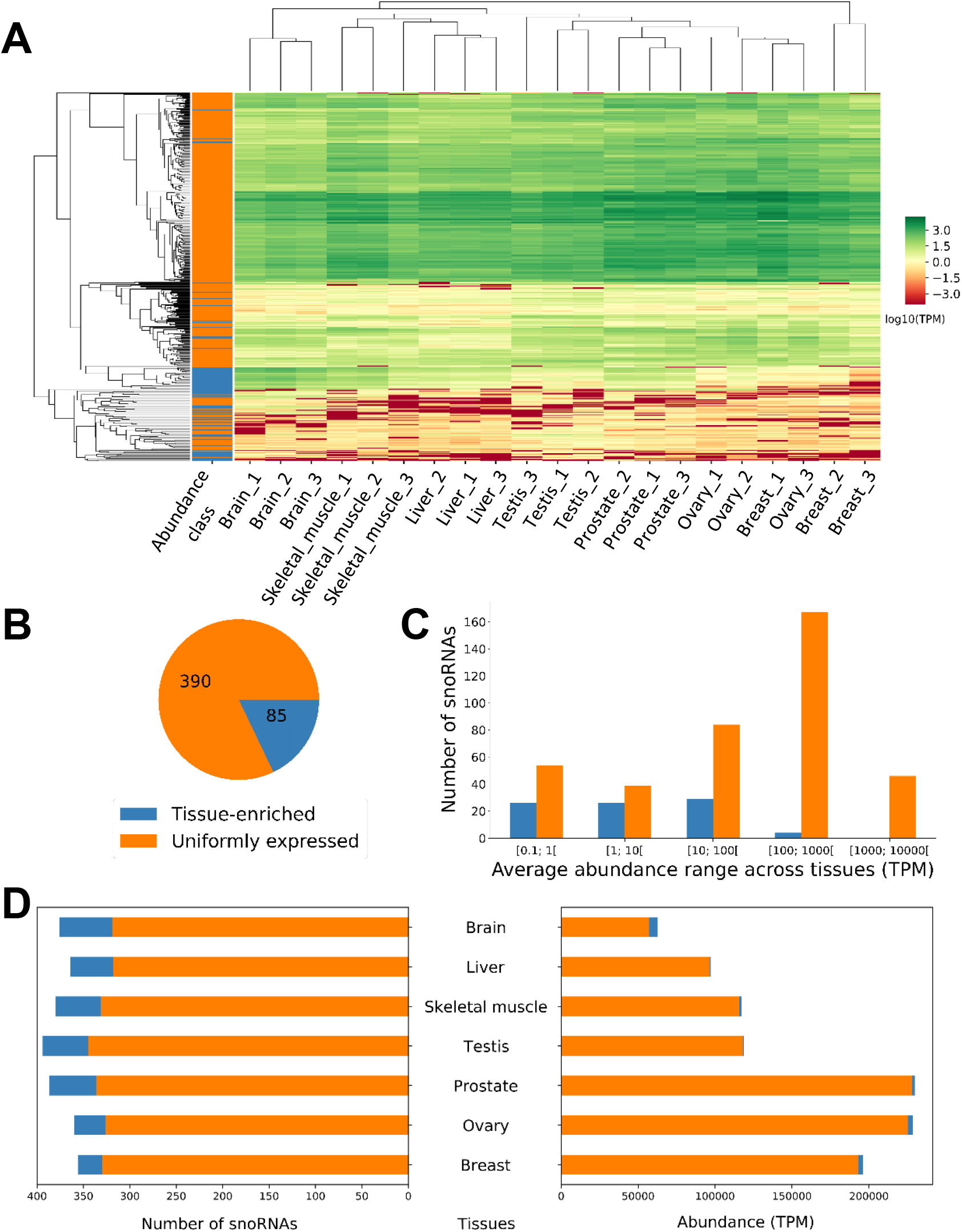
RNA abundance divides snoRNAs into two classes with distinct tissue distribution. (A) SnoRNA abundance accurately clusters healthy human tissues. The heatmap indicates the abundance level of the expressed snoRNAs in 21 RNA samples from 7 different human tissues (legend for the abundance class color bar on the left is shown in (B)). Only snoRNAs with an abundance greater than 1 TPM in at least one sample are considered. The color scale for the abundance is shown on the right. (B) SnoRNAs are divided into two abundance classes. The number of snoRNAs in the uniformly expressed and tissue-enriched classes is shown in the form of a pie chart. (C) Most tissue-enriched snoRNAs are less abundant than their uniformly expressed counterparts. The bar chart indicates the number of both tissue-enriched and uniformly expressed snoRNAs in the different ranges of average RNA abundance across tissues. (D) Most tissue-enriched snoRNAs are expressed in the brain. The number of uniformly expressed and tissue-enriched snoRNAs with an abundance of at least 1 TPM in the examined tissues is displayed on the left, while the total RNA abundance of each class of snoRNAs detected in the different tissues is shown on the right.

Most UE snoRNAs are present at an abundance of >1 TPM in all the examined tissues and the majority has an abundance greater than 100 TPM whereas, in contrast, many TE snoRNAs have an abundance below 1 TPM in most tissues and the majority has an abundance that is less than 100 TPM (Figures S7A, 2C and 2D, left panel). Interestingly, most of TE snoRNA total abundance is attributable to their expression in the brain, whereas UE snoRNA total abundance is mostly attributable to their expression in reproductive tissues (except for testis) (Figure 2D, right panel).

### The snoRNA abundance classes exhibit distinct RNA levels, target preference and conservation patterns

The discovery of two snoRNA abundance classes raises the question of whether the tissue-dependent expression of snoRNAs reflects functional specialization, different evolutionary origin, snoRNA type or simple stochastic variation in expression. To differentiate between these possibilities, we first examined the variation in the abundance of the UE and TE classes in each of the different tissues. As indicated in Figure 3A, all tissues display a broad spectrum of RNA abundance for both groups. Notably, we observe a loose and subtle inverse correlation between the abundance of the two groups: the tissue expressing the lowest amount of TE snoRNAs (Figure 3A, breast tissue) appears to express the highest level of the UE class and vice versa. This suggests that the distribution of these two classes is not random but reflects a tissue-specific expression program that chooses between the housekeeping UE snoRNAs and the specialized TE snoRNAs. To determine whether the abundance classes are driven at least in part by snoRNA type, we then compared the proportion of box C/D and H/ACA snoRNAs in each class. As indicated in Figure 3B, box C/D snoRNAs are well represented in both classes, but the greatest difference is observed with box H/ACA snoRNA, which are significantly more represented in the UE class (Fisher’s exact test, *p*<0.001). These differences in abundance and snoRNA type appear to reflect a degree of functional specialization of the snoRNA abundance classes. Indeed, examining the type of RNA targeted by the snoRNA classes, we notice clear differences in the groups’ target preferences. In general, most targets of the UE class are in rRNA or snRNA, while most TE snoRNAs have no known canonical targets (Fisher’s exact test, *p*<2×10^-14^) (Figure 3C).

**Figure 3.**
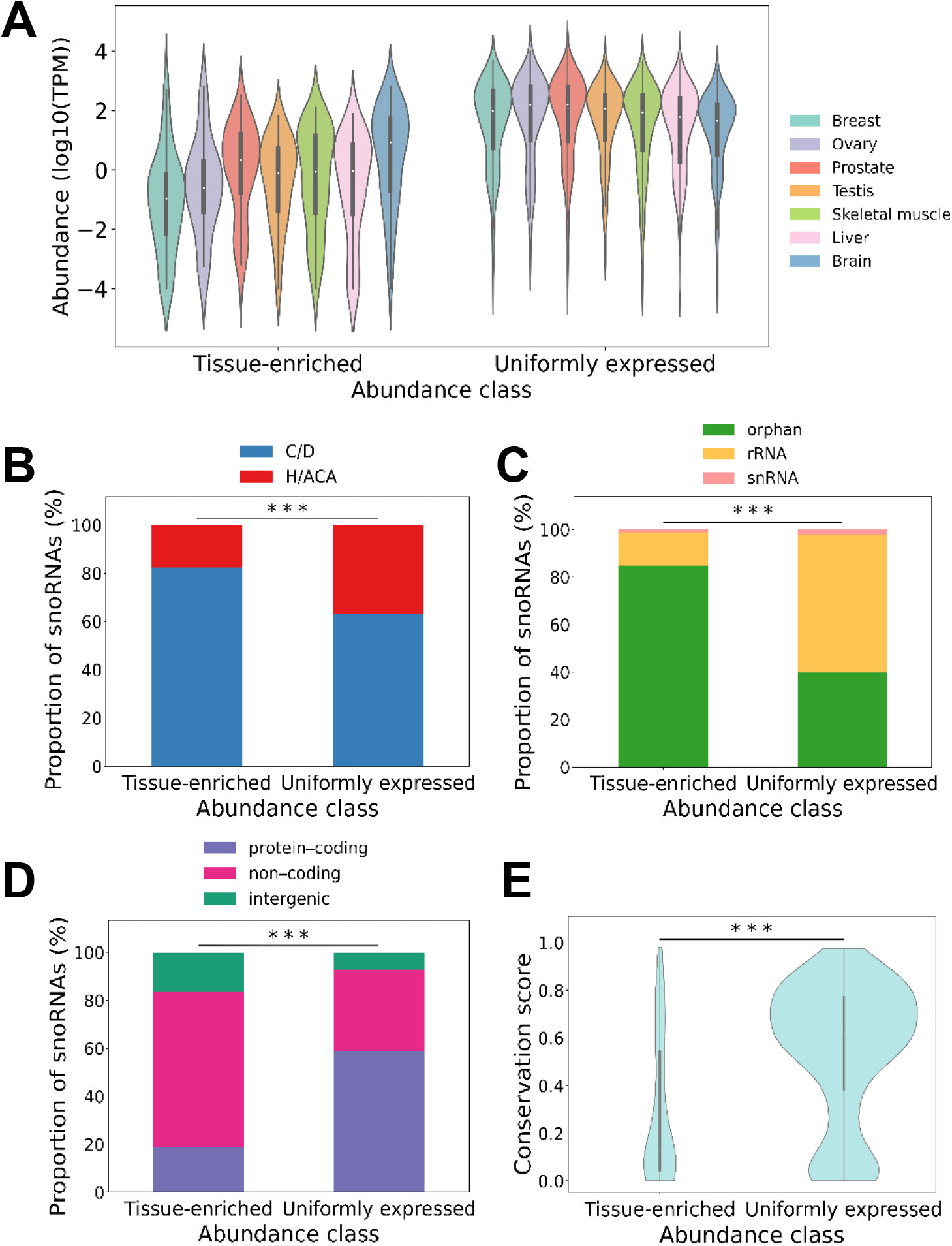
The snoRNA abundance classes represent two groups with distinct characteristics, RNA levels and conservation patterns. (A) The TE and UE snoRNA abundances peak in different tissue types. The abundance of snoRNAs in each tissue per abundance class is represented in the form of a violin plot. (B) Box C/D snoRNAs form the majority of both abundance classes. The percentage of box C/D and H/ACA snoRNAs in each abundance class is shown in the form of a stacked bar chart. The stars indicate the statistical significance of the difference between the two classes of snoRNA abundance (Fisher’s exact test \*\*\**p*<0.001). (C) Most TE snoRNAs are orphans, whereas most UE snoRNAs target rRNA. The stacked bar charts indicate the distribution of the targets (either rRNA, snRNA or orphan, i.e. no known canonical target) of the different snoRNAs in each abundance class. The stars indicate the statistical significance of the difference between the two classes of snoRNA abundance (Fisher’s exact test \*\*\**p*<2×10^-14^) (D) Most TE snoRNAs are embedded in the introns of non-coding HGs, whereas UE snoRNAs are mostly embedded in the introns of protein-coding HGs. The distribution of snoRNA HG biotype in each abundance class is shown as a stacked bar chart. The stars indicate the statistical significance of the difference between the two classes of snoRNA abundance (Fisher’s exact test \*\*\**p*<2×10^-10^). (E) TE snoRNAs are less evolutionarily conserved than the UE class. The violin plots indicate the distribution of the phastCons conservation score in vertebrates for each abundance class. The stars indicate the statistical difference between the two distributions (Mann-Whitney U test, \*\*\**p*<8×10^-11^).

To further characterize the differences between the two snoRNA abundance classes, we compared the genomic organization and conservation of the genes in each class. Interestingly, we found that while the majority of UE snoRNAs are embedded in the introns of protein-coding genes, the majority of the TE snoRNAs are embedded in the introns of non-coding HGs (mainly lncRNAs) (Fisher’s exact test, *p*<2×10^-10^) (Figure 3D). The presence of snoRNAs in non-coding HGs also suggests a more modern evolutionary origin, since many lncRNAs show low sequence conservation [38]. Indeed, comparison of the gene conservation between the two snoRNA groups indicates that the UE class is much more conserved among vertebrates than TE snoRNAs (Mann-Whitney U test, *p*<8×10^-11^) (Figure 3E). TE snoRNAs also tend to be slightly more conserved across primates than vertebrates, but still significantly less than UE snoRNAs (Mann-Whitney U test, *p*<4×10^-9^) (Figures S7B and 3E), highlighting the fact that some TE snoRNAs are potentially only conserved in humans. Altogether, these results indicate that the snoRNA abundance classes represent two groups of snoRNAs with distinct genomic context, conservation, expression patterns and function.

### The snoRNA abundance classes display different degrees of correlation with their HG depending on their HG function and characteristics

Since most snoRNAs in the human genome are embedded in introns [39,40], it is presumed that their expression is linked to that of their HG. To further characterize the relationship between the abundance of snoRNAs and their HG, we thus calculated Pearson correlation coefficients (Pearson’s r or correlation of abundance) and their associated false discovery rate (FDR)-adjusted p-value based on the abundance of the different snoRNA/HG pairs across tissues (Figure 4A). Surprisingly, we find that 40 % of expressed snoRNAs are either non-correlated (−0.25 ≤ correlation of abundance ≤ 0.25) or anticorrelated (correlation of abundance < −0.25) with the abundance of their HG transcripts, suggesting that not all snoRNAs are linked to the expression of their HG and supporting recent findings in other models [22–24]. Indeed, only 60 % of snoRNAs are positively correlated with their HG (correlation of abundance > 0.25) (Figure 4A). The difference in the correlation patterns is not linked to the abundance of snoRNAs as we find that anticorrelated snoRNAs are expressed at similar levels to non-correlated or positively correlated snoRNAs (Figure S8A). On the other hand, snoRNAs are generally more abundant than their HG, and the anticorrelated group in particular is significantly more abundant than their HGs compared to non- or positively correlated snoRNAs (Mann-Whitney U test, *p*<0.05 and *p*<0.0005, respectively) (Figure S9). In addition, we find that in general anticorrelated snoRNAs, regardless of their HG biotype, are more evolutionarily conserved than the other two correlation classes, which underlines their importance in the snoRNome (Figure S8B).

**Figure 4.**
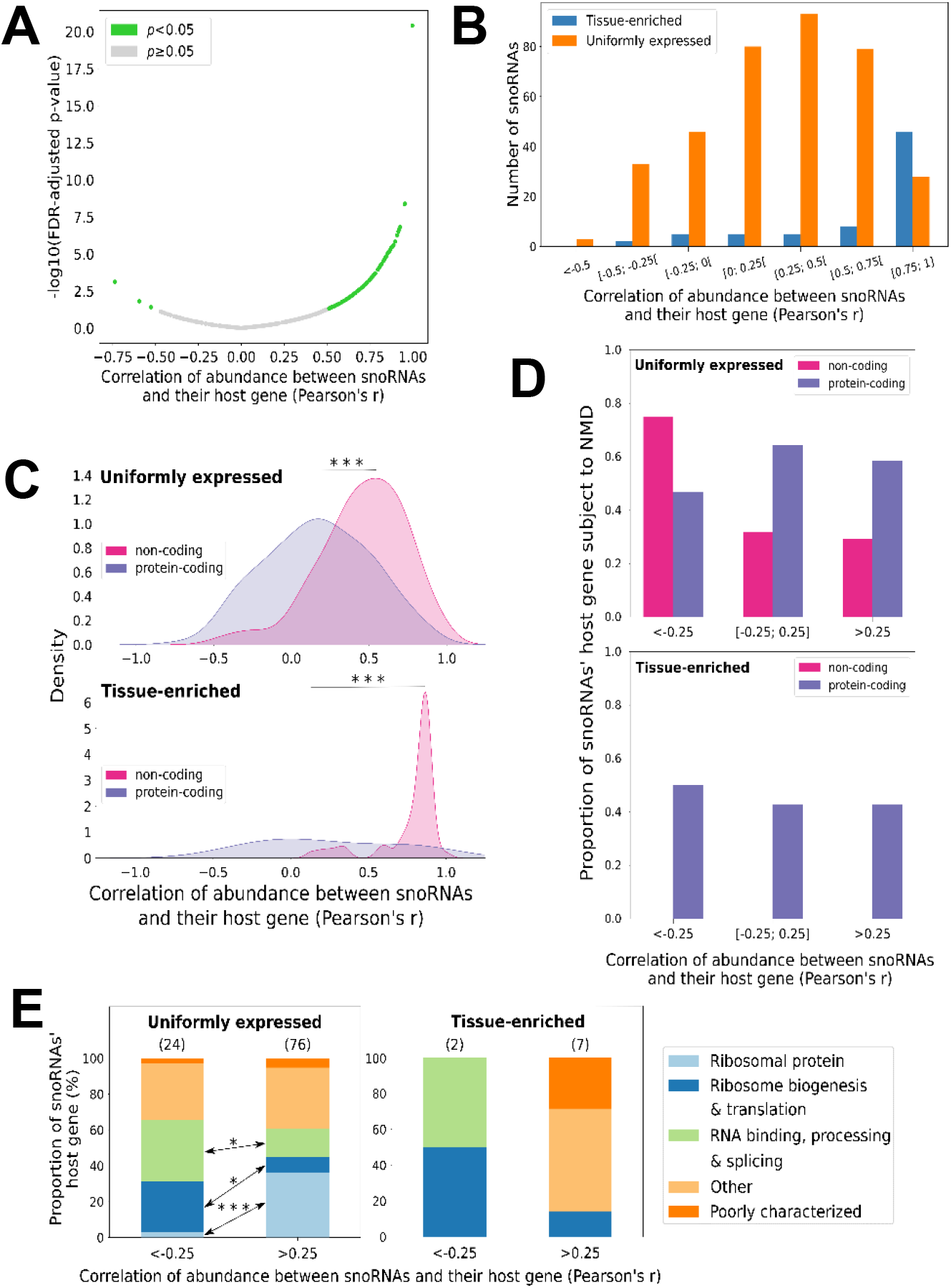
The snoRNA abundance classes correlate differently with their HG abundance due to different HG characteristics. (A) SnoRNAs display a wide range of correlation with their HG abundance. The scatter plot indicates the correlation of abundance of the snoRNA/HG pairs and their associated false discovery rate (FDR)-adjusted p-value for each snoRNA. The green and grey dots indicate respectively significant (*p*<0.05) and non-statistically significant correlations. (B) The abundance of most TE snoRNAs positively correlates with that of their HG as opposed to UE snoRNAs. The number of snoRNAs displaying various degrees of correlation depending on the abundance class is represented as a bar graph. (C) Non-coding HGs are more positively correlated with their embedded snoRNAs than protein-coding HGs. Shown are the density distributions for either UE or TE snoRNAs as a function of the correlation of abundance between the snoRNA and either their protein-coding or non-coding HG. The stars represent the statistical significance of the difference between the two distributions (Mann-Whitney U test, \*\*\**p*<4×10^-15^ and \*\*\**p*<1×10^-5^, respectively for UE and TE snoRNAs). (D) Most anticorrelated non-coding HGs are subject to NMD. The proportion of protein-coding and non-coding HGs subject to NMD is plotted as a function of the correlation of abundance with their embedded snoRNAs (<-0.25: anticorrelation, [-0.25; 0.25]: non-correlation and >0.25: positive correlation), for each snoRNA abundance class. (E) Correlation between UE snoRNAs and their HG abundance is determined at least in part by the HG function. The proportion of anticorrelated and positively correlated snoRNAs embedded in each functional HG group is shown as a stacked bar chart, for each snoRNA abundance class. The number above each bar represents the number of HGs in that subgroup. The stars indicate the statistical significance of the difference between anticorrelated and correlated groups of HGs (Fisher’s exact test, \**p*<0.05 and \*\*\**p*<2×10^-4^).

Since snoRNA abundance spans a wide and variable range of correlation with the HG abundance (Figure 4A), we next wanted to uncover where the two snoRNA abundance classes occur within this broad range of correlation. Interestingly, the TE snoRNAs are much more likely to be correlated with the abundance of their HG transcripts than the UE class, which is represented all along the spectrum of correlation of abundance with the HG (Figure 4B). Since UE and TE snoRNAs have distinct embedding preferences (Figure 3D), we then re-examined the distribution of correlation of abundance, but this time by splitting the two snoRNA abundance classes based on their HG coding potential (Figure 4C). Remarkably, non-coding HGs display clear positive correlation of abundance with either UE or TE snoRNAs, whereas protein-coding HGs exhibit a more complex abundance relationship with their embedded snoRNAs (Mann-Whitney U test, *p*<4×10^-15^ and *p*<1×10^-5^, respectively for UE and TE snoRNAs) (Figure 4C). Overall, these findings suggest that snoRNAs are not always strictly linked to the expression of their HGs and that the snoRNA abundance classes display distinct patterns of correlation with their HG.

Given that snoRNA abundance classes displayed differences in their HG coding potential, we examined the possibility of a link between the snoRNA abundance patterns and the function of their protein-coding genes. Remarkably, we find that UE and positively correlated snoRNAs are predominantly embedded in HGs coding for ribosomal protein (Fisher’s exact test, *p*<2×10^-4^) (Figure 4E, left panel). On the other hand, most anticorrelated UE snoRNAs are located in genes coding for RNA processing and ribosome biogenesis factors (Fisher’s exact test, *p* < 0.05) (Figure 4E, left panel). A similar pattern is observed in the few protein-coding HGs harboring TE snoRNAs, but the small number of HGs prevents accurate estimation of statistical significance (Figure 4E, right panel). Following the same logic but with non-coding HGs, we explored the possibility that lncRNA functionality could be associated with a snoRNA’s correlation of abundance. Indeed, based on previous characterizations of lncRNAs in human diseases [41], those with documented functions are significantly more positively correlated with the abundance of their embedded snoRNAs than lncRNAs with no reported function (Mann-Whitney U test, *p*<2×10^-21^) (Figure S10). Altogether, these results indicate that correlation between the snoRNAs and their HG reflects at least in part the functional relationship of these pairs.

An important characteristic of snoRNA HG groups is the differing stability of their transcripts. Indeed, as reported in [42], mature transcripts encoding ribosomal proteins have a significantly lower decay rate than transcripts from other host gene groups (Mann-Whitney U test, *p*<0.01) (Figure S11A). The abundance of the highly stable mRNA encoding ribosomal protein correlates better in general with their embedded snoRNAs than other HG types (Figure S11B). This increased correlation between the abundance of snoRNA and their host ribosomal protein mRNA may reflect high demand for ribosome synthesis and the need for coordinating the different steps of ribosome biogenesis. We do not observe strong links between the snoRNA-HG correlation of abundance and the stability of non-ribosomal protein host mRNA (Figure S11B). In addition, we note that the stability of the host mRNA will affect the ratio of snoRNA to HG transcript abundance within a tissue, but not their correlation of abundance since the decay rate is presumed to be constant across tissues.

To understand the basis of the difference in the abundance pattern of anti-, non- and positively correlated HGs, we evaluated the susceptibility of HGs to NMD, bearing in mind that NMD could regulate HG transcript levels and thereby modulate the correlation of abundance. NMD-sensitive HGs were defined as such based on their previously determined response to the depletion of NMD factors (see Methods for more details) [21]. Interestingly, we find an increased susceptibility to NMD in anticorrelated non-coding HGs which are enriched in the UE snoRNA class (Figure 4D, top panel). In contrast, we find no association with NMD in the TE class of snoRNAs. This is due to the lack of anticorrelated non-coding HGs of TE snoRNAs and also because non- and positively correlated non-coding HGs of TE snoRNAs are not subject to NMD (Figure 4D, bottom panel), which is consistent with the fact that most TE snoRNAs are highly correlated with the abundance of their non-coding HG transcripts (Figure 4C). Of note, NMD does not seem to modulate alone the correlation of abundance between protein-coding HGs and their embedded snoRNAs, as we observe no significant trend across correlations of abundance for either UE or TE snoRNAs (Figure 4D). Taken together, these findings indicate that NMD may provide means to repress the expression of the HGs without affecting the expression of the embedded snoRNAs and thus enable the uncoupling of the HG and snoRNA expression.

### Dual-initiation of transcription uncouples the expression of the host and snoRNA genes and generates various snoRNA abundance patterns

Since it was recently suggested that promoters with dual transcription initiation sites may uncouple the expression of host and snoRNA genes [25], we compared the number of HGs with only one type of transcription initiation site (termed here simple-initiation sites (SI)) to those with dual-initiation (DI) promoters in both the UE and TE classes of snoRNAs. In addition to a canonical initiation promoter with pyrimidine/purine (YR) dinucleotide, DI promoters carry an additional intertwined polypyrimidine initiation site (YC or 5’TOP) [25]. We therefore defined HGs with DI promoters based on the presence of both YR and YC initiation sites within the HG, which were previously reported using Cap analysis gene expression sequencing (CAGE-Seq) [25] (see Methods for more details). All other HGs were considered as containing an SI promoter. Interestingly, we find that DI promoters are significantly more present in non- and anticorrelated snoRNA/HG pairs regardless of whether they are UE or TE (*p*<6×10^-4^ and *p*<2×10^-5^, respectively for UE and TE snoRNAs) (Figure 5A). Furthermore, significantly more HGs with DI promoters than SI promoters are detected in the UE class of snoRNAs (Fisher’s exact test, *p*<3×10^-8^) (Figure 5B). This is consistent with the increased number of non- and anticorrelated genes detected in the UE class of snoRNAs (Figure 4B) and supports the duality of transcription initiation as a means for uncoupling the HG and snoRNA expression. The initiation pattern dependent uncoupling of either UE and TE snoRNA abundance is also supported by the increased susceptibility of HG transcripts produced from DI promoter to NMD when compared to those generated from a SI promoter (Fisher’s exact test, *p*<2×10^-14^ and *p*<0.01, respectively for UE and TE snoRNAs) (Figure 5C). Strikingly, TE snoRNAs produced from DI and SI promoters have distinct tissue distribution patterns. The SI types are mainly enriched in brain and display positive correlation between the snoRNA and HG, whereas the DI types are highly abundant in breast and ovary tissues and are mostly non- or anticorrelated with their HG (Figure 5D). Collectively, these results indicate that DI promoters present a way for cells to independently optimize the expression of the HG and snoRNA to meet the difference in the functional requirements of human tissues.

**Figure 5.**
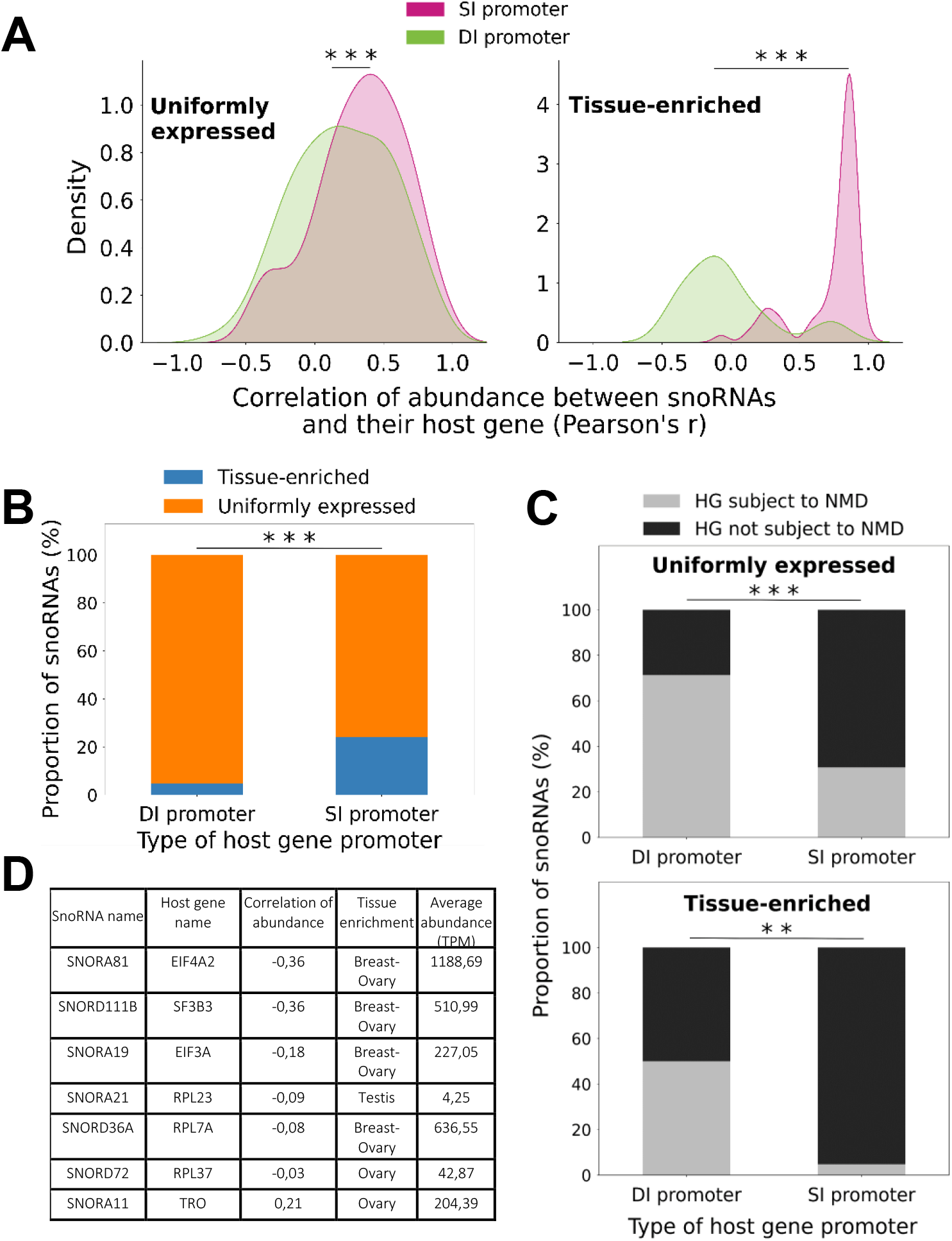
Dual-initiation of transcription uncouples the expression of the host and snoRNA genes and generates various RNA abundance patterns. (A) Most non- or anticorrelated snoRNA/HG pairs are expressed from promoters with dual-initiation sites. The density of snoRNA/HG pairs with dual-initiation (DI promoter) or simple-initiation (SI promoter) promoter is plotted as a factor of the correlation of abundance between snoRNA and their HG, for either UE or TE snoRNAs. The stars indicate the statistical significance of the difference between the DI and SI promoter groups (Mann-Whitney U test, \*\*\**p*<6×10^-4^ and \*\*\**p*<2×10^-5^, respectively for UE and TE snoRNAs). (B) TE snoRNAs are mostly expressed from genes with SI promoters. The proportion of snoRNAs with HGs displaying DI or SI promoters is represented in the form of a stacked bar chart, depending on their abundance class. The stars indicate the statistical significance of the difference between the two HG promoter types (Fisher’s exact test, \*\*\**p*<3×10^-8^). (C) Most HGs displaying DI promoters are subject to NMD. The proportion of snoRNAs with a HG displaying a DI or a SI promoter, depending on if the HG is subject to NMD or not, was plotted as a stacked bar chart, for either UE or TE snoRNAs. The stars indicate the statistical significance of the difference between the two HG promoter types (Fisher’s exact test, \*\*\**p*<2×10^-14^ and \*\**p*<0.01, respectively for UE and TE snoRNAs). (D) Anti- or non-correlated TE snoRNAs embedded in a DI promoter HG are mainly expressed at high levels in female reproductive tissues. The HG name, the correlation of abundance, the tissue enrichment and the average abundance in the enriched tissue(s) of the seven non- or anticorrelated snoRNAs embedded in DI promoter HGs are indicated in a summary table.

## Discussion

In this study, we present a detailed portrait of the human snoRNome and define the basis of snoRNA tissue specificity and abundance patterns. By simultaneously detecting both protein-coding and non-coding RNAs with considerably less structural bias than standard approaches [23,29], we were able to directly compare the snoRNA abundance patterns to the abundance of all non-rRNA biotypes in each studied tissue type (Figure 1), thereby defining a core group of 475 expressed snoRNAs that will serve as valuable resources for future functional analysis. To carry out this analysis, we quantified all human transcripts in seven normal tissues each originating from three different individuals. Interestingly, the data indicate that snoRNAs produce the highest number and diversity of transcripts on average across human tissues after tRNAs (Figure 1). Indeed, unlike snRNAs which occupy a major part of the transcriptome through the expression of only a handful of genes, more than 50 % of snoRNA genes contribute to the abundance of this biotype (Figures S2 and S3). Interestingly, and unlike most highly abundant RNAs in the cells such as snRNAs and tRNAs, not all snoRNAs are uniformly expressed in all tissues (Figures 1 and 2). Instead, a subset of snoRNAs are specifically enriched in brain and reproductive tissues (Figures 2 and S6). Comparison between the UE and TE classes of snoRNAs indicate that they diverge in their target preferences and conservation levels and that the majority of TE snoRNAs are generated from the introns of lncRNAs that mostly correlate with the abundance of their embedded snoRNAs (Figures 3 and 4). In contrast, UE snoRNAs are divided into two groups: the first is highly correlated with its ribosomal protein-coding HG and the second is either non- or anticorrelated with the abundance of its HG transcripts (Figure 4). The non- and anticorrelated snoRNAs are mostly expressed from HGs with DI promoters and their HG transcript is susceptible to NMD, which provides a mechanism to independently regulate the expression of the HG and snoRNA (Figure 5). Overall, the results indicate that snoRNAs are not a mere group of uniformly expressed genes that obey the instruction of their HG but include subgroups with distinct gene organization and abundance patterns that meet the demand for both housekeeping and tissue-specific functions.

Altogether, our data suggest a model in which intron-embedded snoRNA expression patterns and tissue specificity are products of the HG function and architecture (Figure 6). In this model, the majority of TE snoRNAs are encoded in the introns of lncRNA genes, while the majority of UE snoRNAs are encoded in protein-coding genes. Non-coding HGs free the cell to optimize the expression and/or rapidly evolve specialized snoRNAs to meet tissue-specific requirements while embedding snoRNAs within protein-coding genes provides a broad range of regulatory relationships between the snoRNA and host protein functions. Indeed, the majority of non-coding HGs use uncomplicated expression modules where the abundance of the host transcript and snoRNA are positively correlated (Figure 6, third expression module from the left), whereas in contrast, most protein-coding HGs are non- or anticorrelated with the abundance of their embedded snoRNAs (Figure 6, second expression module from the left).

**Figure 6.**
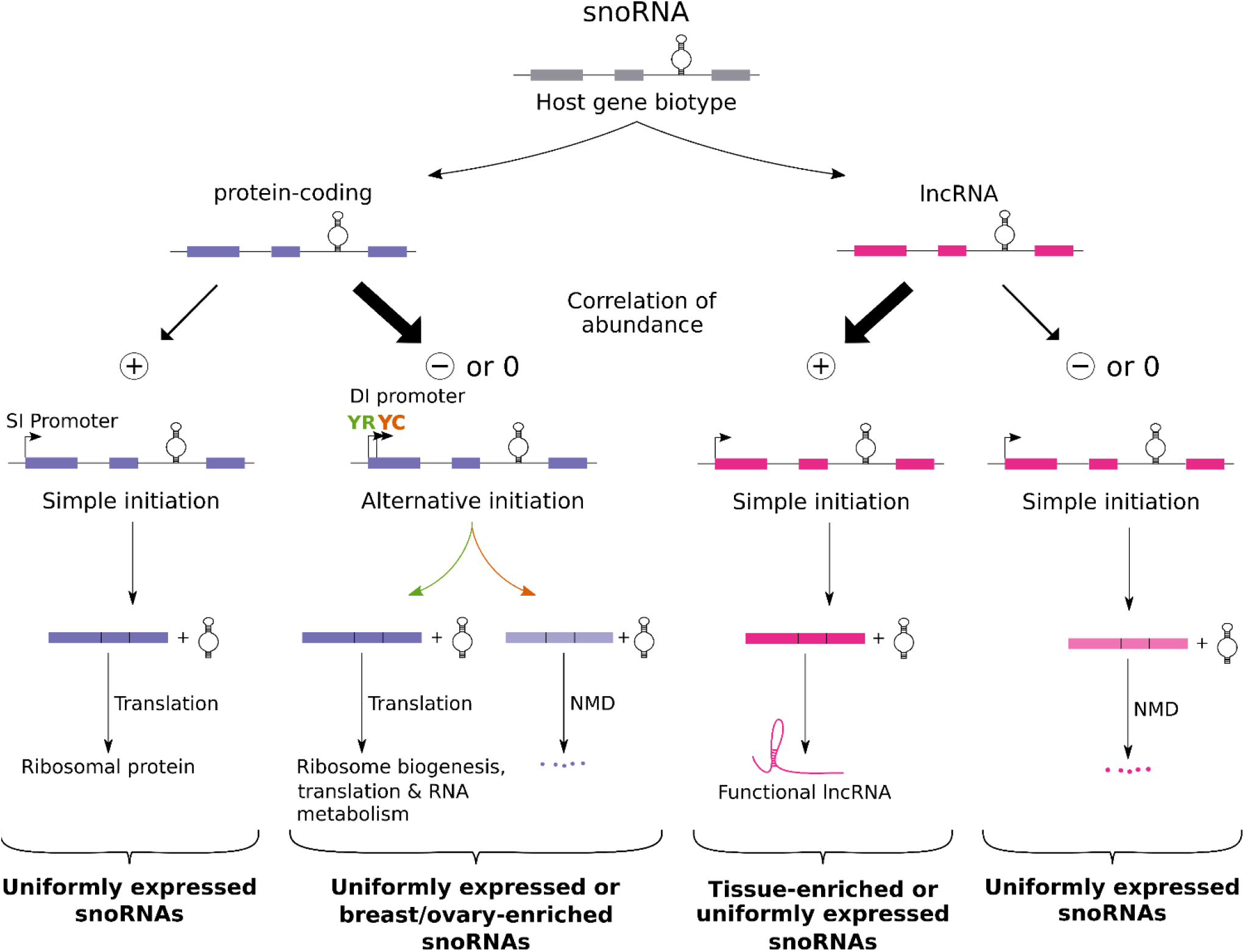
Model explaining the modulation of snoRNA expression. In this model, the abundance pattern of snoRNAs is determined by the host function and transcription pattern. Alternative initiation of transcription (either canonical YR-initiation or non-canonical YC-initiation) provides a means to independently regulate the expression of the snoRNA and its HG and is often found in UE and a few TE snoRNAs embedded in protein-coding HGs. On the other hand, simple transcription initiation is the hallmark of positively correlated snoRNA/HG pairs. Only the main pathways and features determining the snoRNA abundance patterns are illustrated. The anticorrelated, non-correlated and positively correlated snoRNAs are indicated by -, 0 and +, respectively. Bold arrows represent the principal expression pathways adopted by snoRNAs. NMD, SI and DI promoters stand for nonsense-mediated decay, simple-initiation and dual-initiation promoters respectively.

Almost all positively correlated snoRNAs are expressed from promoters with a simple transcription initiation site, confirming their obligate joint expression pattern (Figure 6, first and third expression modules from the left). Conversely, non- or anticorrelated snoRNAs embedded in lncRNAs are also generated through SI transcription, but since the HG transcript has no known associated function and is highly susceptible to NMD, only a stable UE snoRNA remains after the transcription of the HG that thereby serves the only purpose of expressing its embedded snoRNA (Figure 6, right-most expression module). Given the correlated expression pattern of host and snoRNA genes combined with the observed insensitivity to NMD of positively correlated snoRNA-containing lncRNAs (Figure 6, third expression module from the left), it is thus likely that these stable lncRNAs play compatible or complementary roles with their embedded snoRNAs. Interestingly, most brain-enriched snoRNAs, which are encoded in the Prader-Willi syndrome region, are generated through this and joint expression with their non-coding HG (Figure 6, third expression module from the left). Genes of this genomic region were recently reported to produce 5’-snoRNA-capped and 3’-polyadenylated lncRNAs (SPAs) and lncRNAs flanked by snoRNA (sno-lncRNAs), which are hybrids involved in RNA binding protein trapping [18,43,44]. This suggests that these TE snoRNA/lncRNAs pairs either work as a whole or as separate entities to achieve common tissue-specific functions. Following the same logic, positively correlated protein-coding HGs (Figure 6, first expression module from the left) produce through simple expression both the snoRNA and the HG transcript, which is most likely coding for a ribosomal protein. Since this expression module produces UE snoRNAs, which mostly target rRNA, this underlines that a positive correlation of abundance reflects a functional link between snoRNAs and their HG: UE snoRNA-guided modification of rRNA and ribosomal proteins being both important factors of ribosome structure integrity [45].

In contrast to the simple positive expression module of most TE snoRNAs, the majority of UE snoRNAs and few ovary- and breast-enriched snoRNAs use a complex regulatory module that separates the expression of the snoRNA from its HG (Figure 6, second expression from the left). In most cases, this separation of expression is achieved through DI promoters that use different transcription initiation sites (either canonical YR-initiation or non-canonical YC-initiation) depending on the need of the different tissues. The snoRNA is expressed regardless of the initiation site, but the host transcript accumulates only when the YR-initiation is used, which protects the transcript from degradation by NMD. In this way, the cell may regulate the expression of the HG without interfering with the uniformity of snoRNA abundance, which likely responds to the need for snoRNAs with a housekeeping function such as most UE snoRNAs. As expected, the non- and anticorrelated HGs using DI sites are not enriched in housekeeping genes like ribosomal protein genes. Instead, they mainly include genes that regulate RNA maturation and processing such as genes involved in ribosome biogenesis. Indeed, it seems that in most cases the separation of HG and snoRNA functions is needed to liberate the snoRNA from tissue and condition dependent control of the HG. Interestingly, in few cases like SNORD63 and SNORD50A, the promoter duality may even allow the snoRNA to develop non-canonical functions such as regulating pre-mRNA stability and polyadenylation [46,47]. Further studies are however needed to characterize the biological relevance of a lack of positive correlation between a snoRNA and its HG and to decipher what distinguishes anticorrelated from non-correlated snoRNAs. Collectively, the data presented here and summarized in Figure 6 indicate that the human snoRNome meets the demands of both uniform and tissue-enriched abundance through a broad spectrum of regulatory mechanisms that define the relationship between the snoRNA and its HG expression.

## Conclusions

SnoRNAs are implicated in a myriad of crucial functions in eukaryotic cells, yet their abundance patterns across healthy human tissues and their relationships with their HG had never been comprehensively studied. In this study, we generated fragmented and ribodepleted TGIRT-Seq abundance datasets of both structured and non-structured RNAs in seven healthy human tissues, enabling us to reliably characterize for the first time the entire human snoRNome. SnoRNAs were identified as major contributors of the abundance in all the tissues and were divisible in two abundance classes with clear and distinct characteristics: UE and TE snoRNAs. Almost half of all expressed snoRNAs were found to be non- or anticorrelated with the abundance of their HG transcripts, highlighting a complex abundance regulation. The HG function and promoter duality were identified as crucial features that modulate the abundance patterns of snoRNAs and their HG in order to meet the functional requirements of both UE and TE snoRNAs in human tissues. Overall, our study represents a reliable reference from which future research can draw upon to better characterize the importance of snoRNAs in human physiological and pathological conditions.

## Methods

### Sample origin and preparation

RNA from healthy skeletal muscle, liver, testis and brain tissues was purchased from BioChain (3 RNA samples per tissue originating from different individual donors). Healthy breast, ovary and prostate tissue samples were obtained from the FRSQ tissue bank (Université de Sherbrooke). Each 30 mg tissue sample was homogenized in 1 mL of TRIzol Reagent (Ambion) using a Polytron tissue homogenizer and kept at −80°C until RNA extraction. Characteristics of the samples are available in Table S4.

### RNA extraction

Since RNA was directly purchased for the skeletal muscle, liver, testis and brain tissues, only total RNA extractions from breast, ovary and prostate tissues were performed using RNeasy Mini Kit (Qiagen) as recommended by the manufacturer including on column DNase digestion with RNase-Free DNase Set (Qiagen). However, 1.5 volume of ethanol 100 % was used instead of the recommended 1 volume of ethanol 70 % in order to retain smaller RNA. RNA integrity of each sample was assessed with a 2100 Bioanalyzer (Agilent). These values are available from Table S4 for all samples.

### Ribodepletion, library preparation and paired-end sequencing

RNA-Seq libraries were built as previously described [23]. Briefly, 2 μg of DNA-free total RNA was ribodepleted using Ribo-Zero Gold (Illumina) according to the manufacturer protocol. The resulting rRNA-depleted RNA was then purified with RNA Clean and Concentrator (RCC) kit (Zymo Research) using a modified protocol to retain RNA ≤ 80 nucleotides (400 μL ethanol 100 % per 50 μL sample). Purified RNA was fragmented 2-4 minutes (depending on the RNA Integrity Number) using NebNext Magnesium RNA Fragmentation Module (New England Biolabs) and once again purified with the RCC kit (Zymo Research) followed by dephosphorylation using T4 Polynucleotide Kinase (Epicentre) and final purification using, again, the RCC kit (Zymo Research).

cDNAs were synthesized via TGIRT template-switching with 1 μM TGIRT-III reverse transcriptase (Ingex, LLC) for 15 min at 60°C, during which a DNA oligonucleotide containing the complement of an Illumina Read 2 sequencing primer-binding site became seamlessly linked to the 5’ cDNA end. After reaction cleanup, a 5’ adenylated DNA oligonucleotide containing the complement of an Illumina Read 1 sequencing primer-binding site was then ligated to the 3’ cDNA end with Thermostable 5’ AppDNA / RNA Ligase (New England Biolabs). Properly ligated cDNAs were amplified by PCR (12 cycles) to synthesize the second strand and add Illumina flowcell capture and index sequences. Libraries were purified with 2 rounds of Ampure XP beads (Beckman-Coulter) and evaluated on a 2100 Bioanalyzer (Agilent). Libraries were then pooled and sequenced on a NextSeq 500 platform (Illumina) (2 x 75 bp) using a NextSeq 500/550 High Output Kit v2.5 (150 cycles) (Illumina). Three distinct sequencing runs were performed to sequence all tissue samples: the first pool was composed of the Breast_1, Breast_2, Ovary_1, Ovary_2, Ovary_3, Prostate_1, Prostate_2 and Prostate_3 RNA samples; the second pool was composed of the Brain_1, Brain_2, Brain_3, Liver_1, Liver_2, Liver_3, Testis_1 and Testis_2 RNA samples; the third pool was composed of the Breast_3, Skeletal_muscle_1, Skeletal_muscle_2, Skeletal_muscle_3 and Testis_3 RNA samples.

### TGIRT-Seq processing pipeline

All RNA abundance datasets were generated using a succession of bioinformatics tools regrouped in a reproducible Snakemake workflow [48]. All details about parameters and tools used can be found in the Snakemake workflow at http://gitlabscottgroup.med.usherbrooke.ca/etienne/tgirt_seq_pipeline, but are also briefly described below. The datasets we generated are of high depth and quality for each tissue (Table S3) and are available for download from the Gene Expression Omnibus (the breast, ovary and prostate datasets are available under the accession number GSE126797 and the remaining datasets are available under the accession number GSE157846). In short, paired-end reads were first trimmed using Trimmomatic v0.36 [49] (with the following parameters: ILLUMINACLIP:<fastaWithAdaptersEtc>:2:12:10:8, TRAILING:30, LEADING:30, MINLEN:20, all other parameters at default values) to remove adapters and low-quality reads. FastQC v0.11.5 was used before and after trimming to assess the quality of the reads. Trimmed reads were aligned to the human genome assembly GRCh38 (hg38, v87) using the aligner STAR v2.6.1a [50] (with the following parameters: --runMode alignReads, --outSAMunmapped None, --outSAMtype BAM SortedByCoordinates, --outFilterScoreMinOverLread 0.3, --outFilterMatchNminOverLread 0.3, --outFilterMultimapNmax 100, --winAnchorMultimapNmax 100, --alignEndsProtrude 5 ConcordantPair, all other parameters at default values). The index needed to align reads to the human genome was generated using STAR v2.6.1a [50] (with the following parameters: --runMode genomeGenerate and --sjdbOverhang 74). Counts were attributed to genomic features using CoCo v0.2.1p4 [33] (with the following parameters: cc -countType both -strand 1 --paired, all other parameters at default values), using our custom annotation (.gtf file available at https://zenodo.org/record/3981426/files/human_ensembl_87_wo_dup_v2.BB_v3.correct_annotation.gtf) described in [23]. Normalized counts in TPM were obtained from the output of CoCo. Only snoRNAs with an abundance greater than 1 TPM in at least one tissue sample, thus referred to as “expressed snoRNAs”, were included in this study in order to filter out low abundance snoRNAs. Also, even though their associated biotype was “snoRNA”, 4 snoRNAs with a gene name starting with “SCARNA” were manually excluded from this analysis.

### Collection of GTEx expression data

The 10 most abundant protein-coding genes in GTEx for the seven tissues studied were manually curated through the GTEx portal [34]. Mitochondrial genes were excluded from both the GTEx and TGIRT-Seq rankings.

### Grouping of RNA biotypes

In order to simplify the analysis, RNA biotypes obtained from our custom annotation were grouped in classes according to Ensembl nomenclature. Thus, IG_C_gene, IG_D_gene, IG_J_gene, IG_V_gene, TR_C_gene, TR_D_gene, TR_J_gene, TR_V_gene, polymorphic_pseudogene and protein_coding biotypes were grouped under the generic “protein-coding” biotype; unitary_pseudogene, unprocessed_pseudogene, processed_pseudogene, transcribed_unprocessed_pseudogene, transcribed_unitary_pseudogene, transcribed_processed_pseudogene, IG_pseudogene, IG_C_pseudogene, IG_J_pseudogene, IG_V_pseudogene, TR_J_pseudogene, TR_V_pseudogene and pseudogene biotypes were grouped under the generic “pseudogene” biotype; 3prime_overlapping_ncRNA, antisense, lincRNA, macro_lncRNA, bidirectional_promoter_lncRNA, processed_transcript, sense_intronic, sense_overlapping, non_coding and lncRNA biotypes were grouped under the generic “lncRNA” biotype; Mt_tRNA and tRNA biotypes were grouped under the generic “tRNA” biotype; rRNA, Mt_rRNA, ribozyme, scRNA, vaultRNA and sRNA biotypes were grouped under the generic “other” biotype. Of note, RNAs with missing abundance value in any tissue sample and RNAs with the “TEC” biotype were not considered in this study. Following the same logic, HG biotypes were grouped under three generic biotypes: “protein-coding” for all protein-coding HG, “intergenic” for snoRNAs without a HG and “non-coding” for all other HG biotypes.

### Collection of snoRNA related information

Protein-coding HG biological functions were manually curated from UniProt [51] and non-coding HG (lncRNAs) associated functions in various human diseases were retrieved from LncTarD [41]. NMD susceptibility of the HG was based on the presence of the HG in the Supplementary table S4 of Lykke-Andersen *et al*. (corrigendum version of the original paper) using their relaxed criterion [21]. This table lists all genes determined as NMD substrates based on their increased accumulation after different depletions of NMD factors [21]. The presence of DI promoters within a HG was defined by the presence of the HG in the Supplementary Data 7 of Nepal *et al*. [25]. This table lists all genes identified as containing both YR and YC promoters using CAGE-Seq in the human HepG2 cell line [25]. The score of conservation across vertebrates (“phastCons 100 Vertebrates”) and across primates (“phastCons 30 primates”) for each snoRNA was obtained from the UCSC Genome Browser [52,53]. In short, a conservation score was associated to each nucleotide of a snoRNA and the conservation score per snoRNA was generated by calculating the average score of all the nucleotides included in that snoRNA sequence. Otherwise, all other information (e.g. a snoRNA’s target, HG name and biotype, etc.) was retrieved from snoDB [40]. SnoRNAs without known target in rRNA or snRNA were designated as “orphan” snoRNAs. All snoRNA abundance and features are available in Table S5.

### Abundance class categorization

To categorize snoRNAs according to their abundance patterns across healthy human tissues, a coefficient of variation (CV) was calculated for each snoRNA. This method was also applied to other RNA biotypes (snRNA, tRNA, protein-coding RNA and lncRNA). In short, the CV was calculated as the standard deviation of the abundance of that snoRNA across the tissues divided by the average abundance of that snoRNA across the tissues, all of that multiplied by 100. These CVs were represented in a kernel density estimate plot and the resulting bimodal curve was divided in two by tracing the tangent at the point where the derivative of the bimodal curve function was the most negative. The point at which the tangent crossed the x-axis was defined as the threshold for the two snoRNA abundance classes. Above that threshold of CV=125, snoRNAs were dubbed “Tissue-enriched” or “TE”, whereas snoRNAs with a CV below that threshold were dubbed “Uniformly expressed” or “UE”. To classify in which tissue TE snoRNAs were predominantly expressed, the tissue where the snoRNA abundance (in TPM) was the highest was established as the enrichment tissue. This was the case for all TE snoRNAs except for 4 snoRNAs (SNORA81, SNORA19, SNORD36A and SNORD111B) that were highly abundant in both breast and ovary and had a difference of abundance (in TPM) of at most 2 times the abundance seen in the other tissue (either breast or ovary).

### Statistical analyses and graph generation

All statistical analyses and graphs were realized using Python-based packages. Pearson correlation coefficients (Pearson’s r) and their associated p-values, Fisher’s exact test p-values and Mann-Whitney U test p-values were generated using the Stats module from Scipy v1.4.1. SnoRNAs with a correlation of abundance with their HG (Pearson’s r) inferior to −0.25 were considered “anticorrelated” whereas those with a Pearson’s r greater than 0.25 were considered “positively correlated”; snoRNAs with a Pearson’s r comprised inclusively between −0.25 and 0.25 were considered “non-correlated”. P-value correction for false-discovery rate (FDR) using the Benjamini-Hochberg correction (for the correlation of abundance between snoRNAs and their HG) was performed using the Multitest module from Statsmodels v0.11.0. Throughout this study, all results were considered significant at \**p* < 0.05, *\*\*p* < 0.01 and *\*\*\*p* < 0.001. Graphs were generated using either the pandas v1.0.1, Matplotlib v3.1.1 or Seaborn v0.9.0 libraries.

## Declarations

### Ethics approval and consent to participate

Not applicable.

### Consent for publication

Not applicable.

### Availability of data and materials

The datasets we generated are available for download from the Gene Expression Omnibus (GEO) repository. The breast, ovary and prostate datasets are available under the accession number GSE126797 (https://www.ncbi.nlm.nih.gov/geo/query/acc.cgi?acc=GSE126797) and the remaining datasets are available under the accession number GSE157846 (https://www.ncbi.nlm.nih.gov/geo/query/acc.cgi?acc=GSE157846). All abundance datasets were generated using a succession of bioinformatics tools regrouped in a reproducible Snakemake workflow that is accessible for download (http://gitlabscottgroup.med.usherbrooke.ca/etienne/tgirt_seq_pipeline).

### Author’s contribution

M.S.S. and S.A.E. conceived the study and designed the experiments. S.C. prepared the samples and the libraries and carried out the sequencing. É.F.C. analyzed and interpreted the data with the help of M.S.S. D.B. provided general wisdom throughout the whole project. É.F.C. and M.S.S. wrote the manuscript and all other authors provided feedback on the manuscript. All authors read and approved the final manuscript.

## Acknowledgments

The authors would like to thank members of the Scott and Abou-Elela groups for helpful discussions and Compute Canada for providing state-of-the-art computing infrastructures.

