## Supplementary Figures and Tables S1-S4 for "Annotation of snoRNA abundance across human tissues reveals complex snoRNA-host gene relationships"

### Additional file 1

**The file includes:**

Figures S1-S11

Tables S1-S4

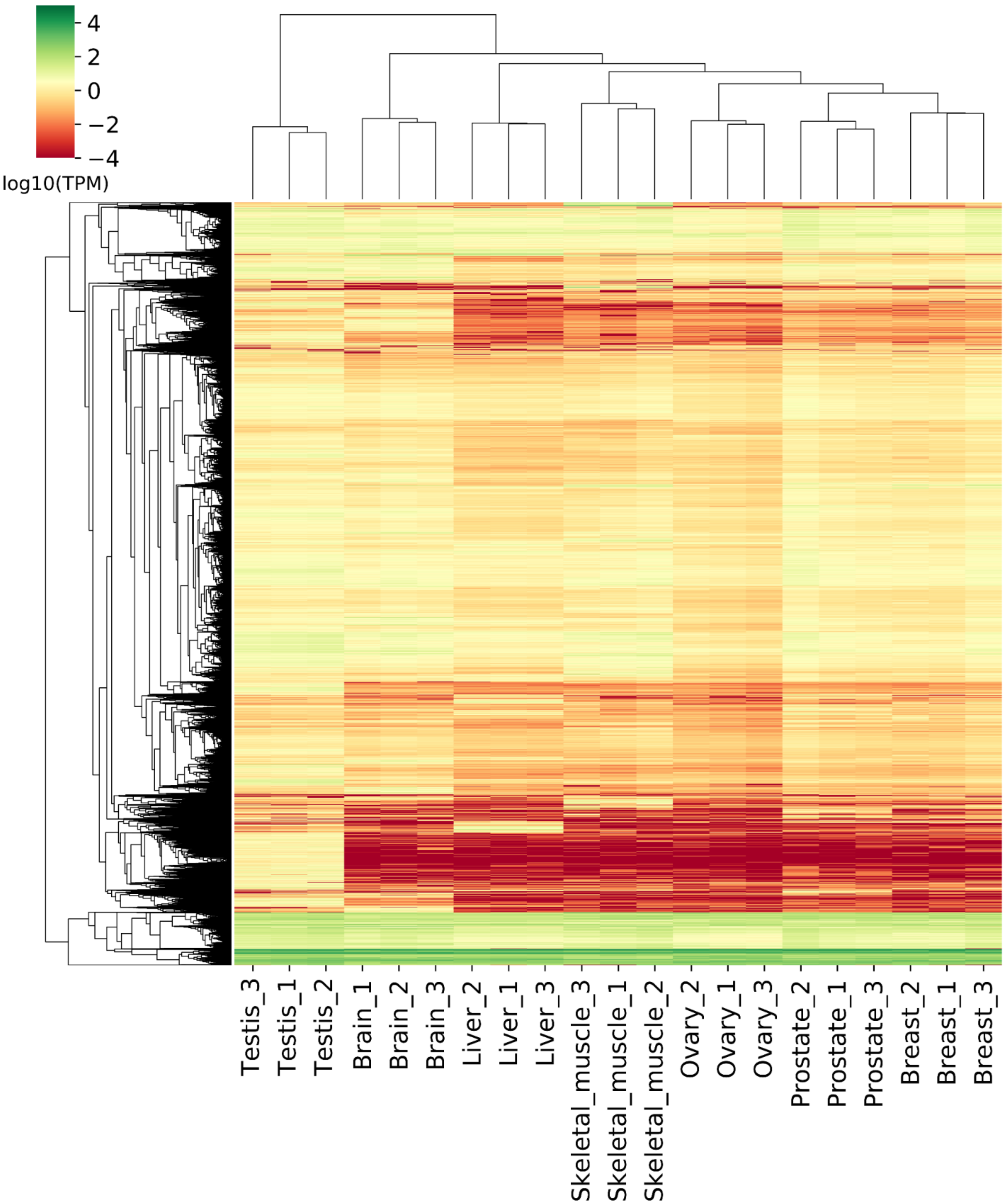


**Figure S1 (Supplementary to Figure 1)**. **Most RNAs are poorly expressed across the tissues.** Heatmap of all RNAs with an abundance greater than 1 TPM in at least one tissue sample. Abundance color scale is shown on top left.

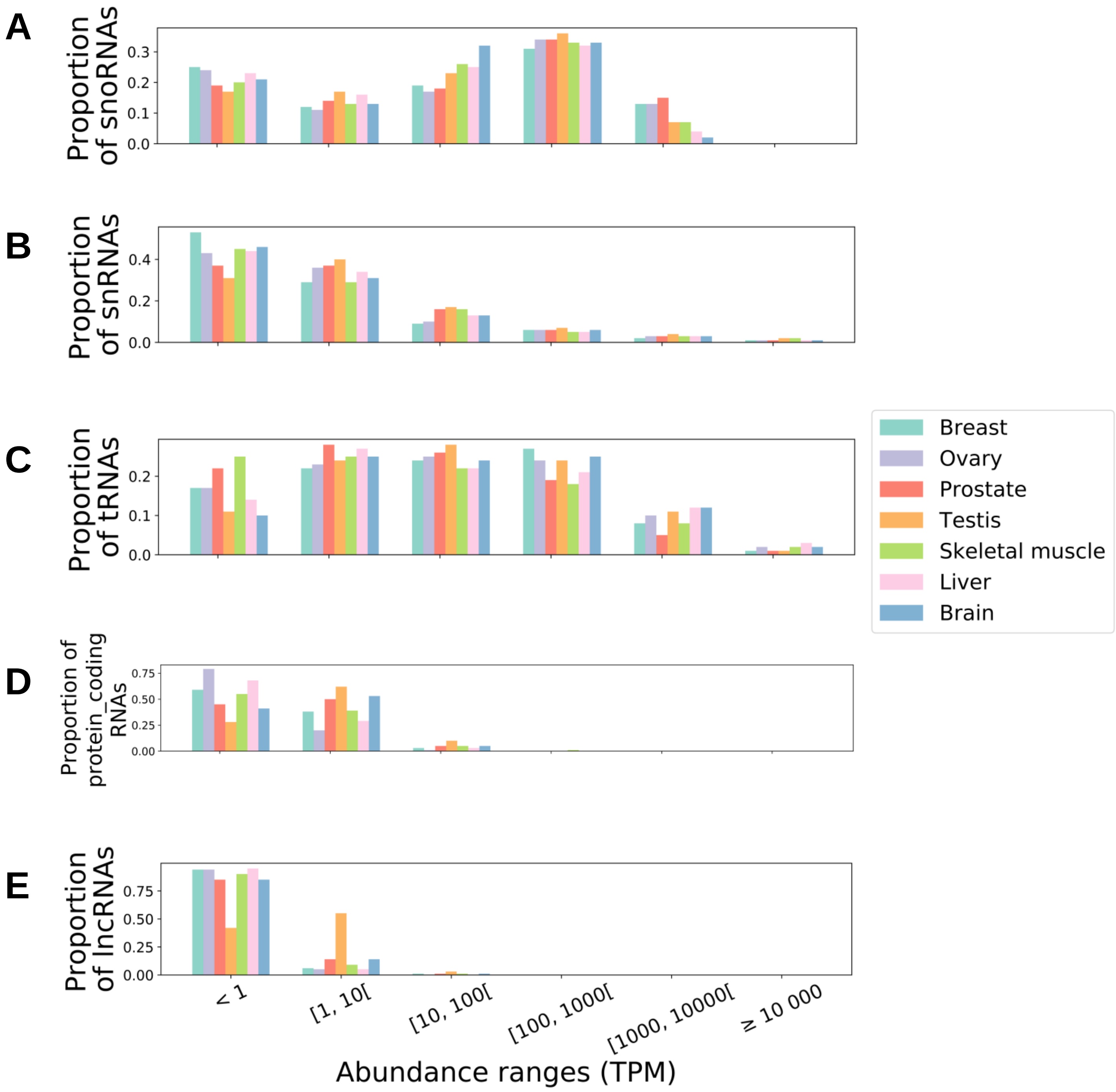


**Figure S2 (Supplementary to Figure 1). SnoRNAs and tRNAs show a wide range of abundance across tissues compared to other biotypes.** (A-E) Bar chart displaying the proportion of a specific RNA biotype across abundance ranges for each tissue for the snoRNA (A), snRNA (B), tRNA (C), protein-coding (D) and lncRNA (E) biotypes.

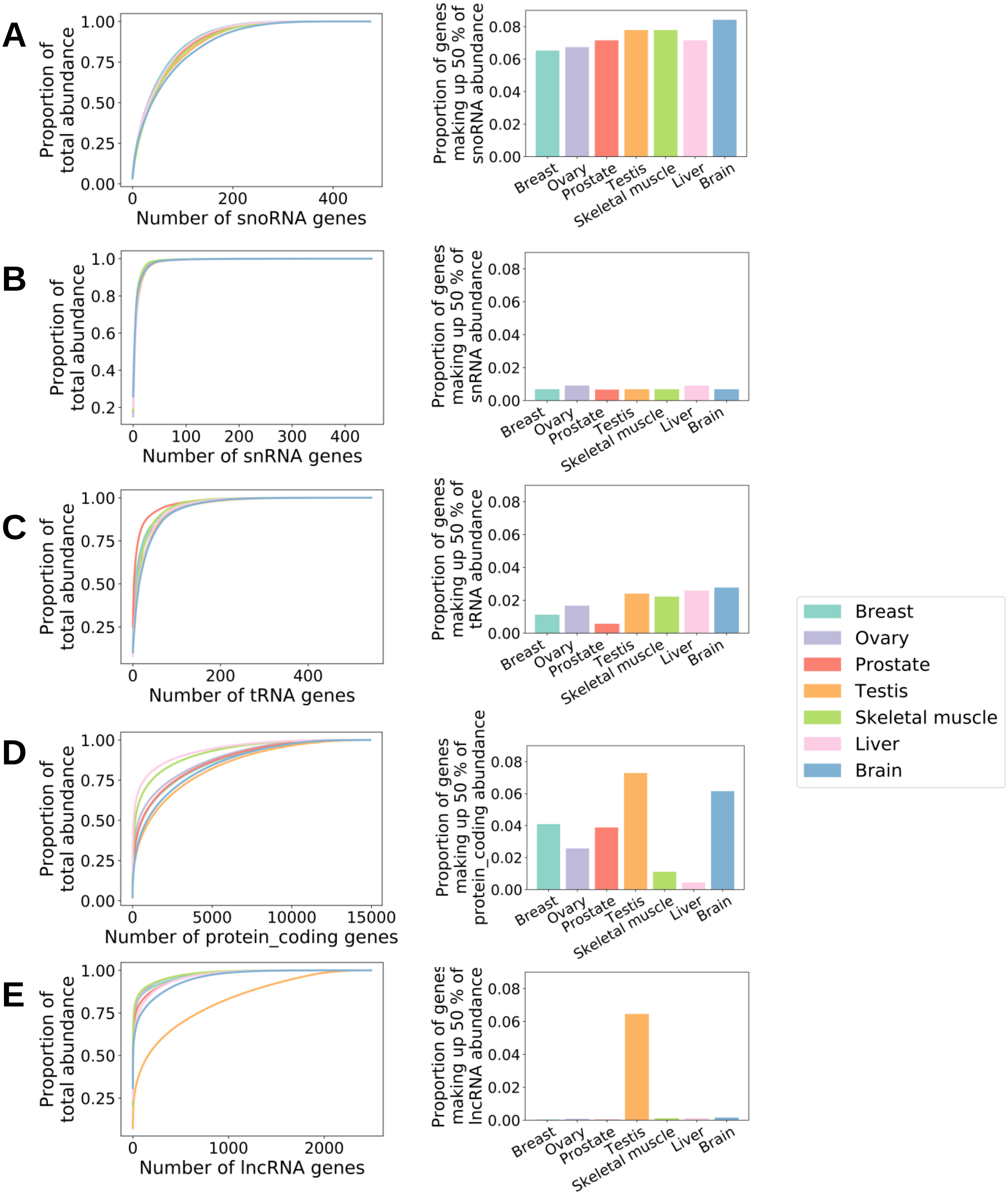


**Figure S3 (Supplementary to Figure 1). SnoRNAs show less variability than protein-coding RNA in their abundance across tissues.** (A-E) Cumulative abundance of gene biotypes expressed at an abundance greater than 1 TPM per tissue (left panel) (legend id shown in far middle right) and bar graph showing the proportion of genes making up the top 50% of the biotype in terms of abundance (right panel) for the snoRNA (A), snRNA (B), tRNA (C), protein-coding (D) and lncRNA (E) gene families.

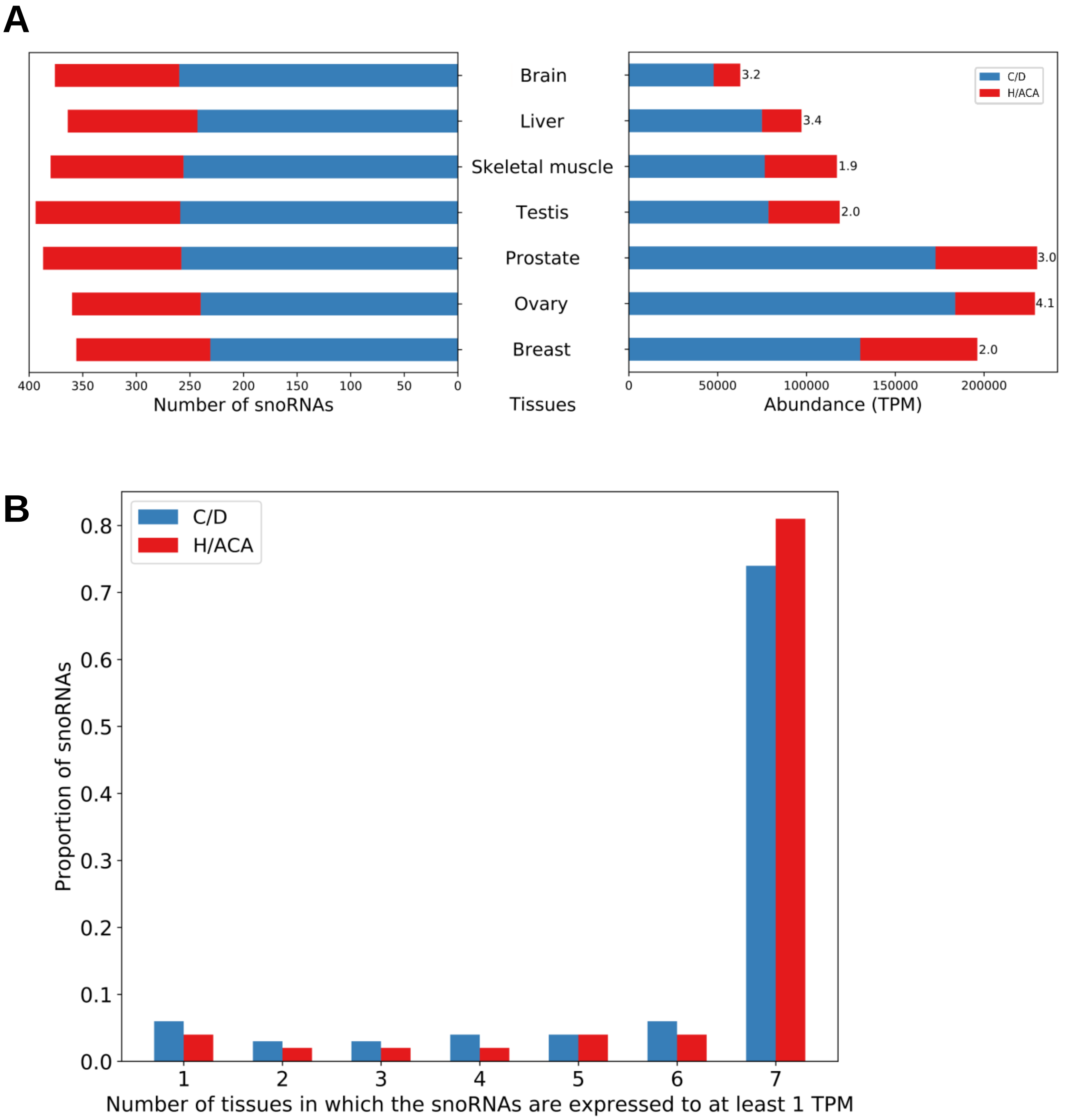


**Figure S4 (Supplementary to Figure 1). Box C/D snoRNAs are more abundant than box H/ACA snoRNAs across tissues.** (A) Number of box C/D and H/ACA snoRNAs that have an abundance greater than 1 TPM across tissues and total abundance of snoRNAs across tissues (box C/D to H/ACA abundance ratio shown on the right of the bars). (B) Proportion of snoRNAs of type box C/D or H/ACA as a function of the number of tissues in which they are abundant to at least 1 TPM.

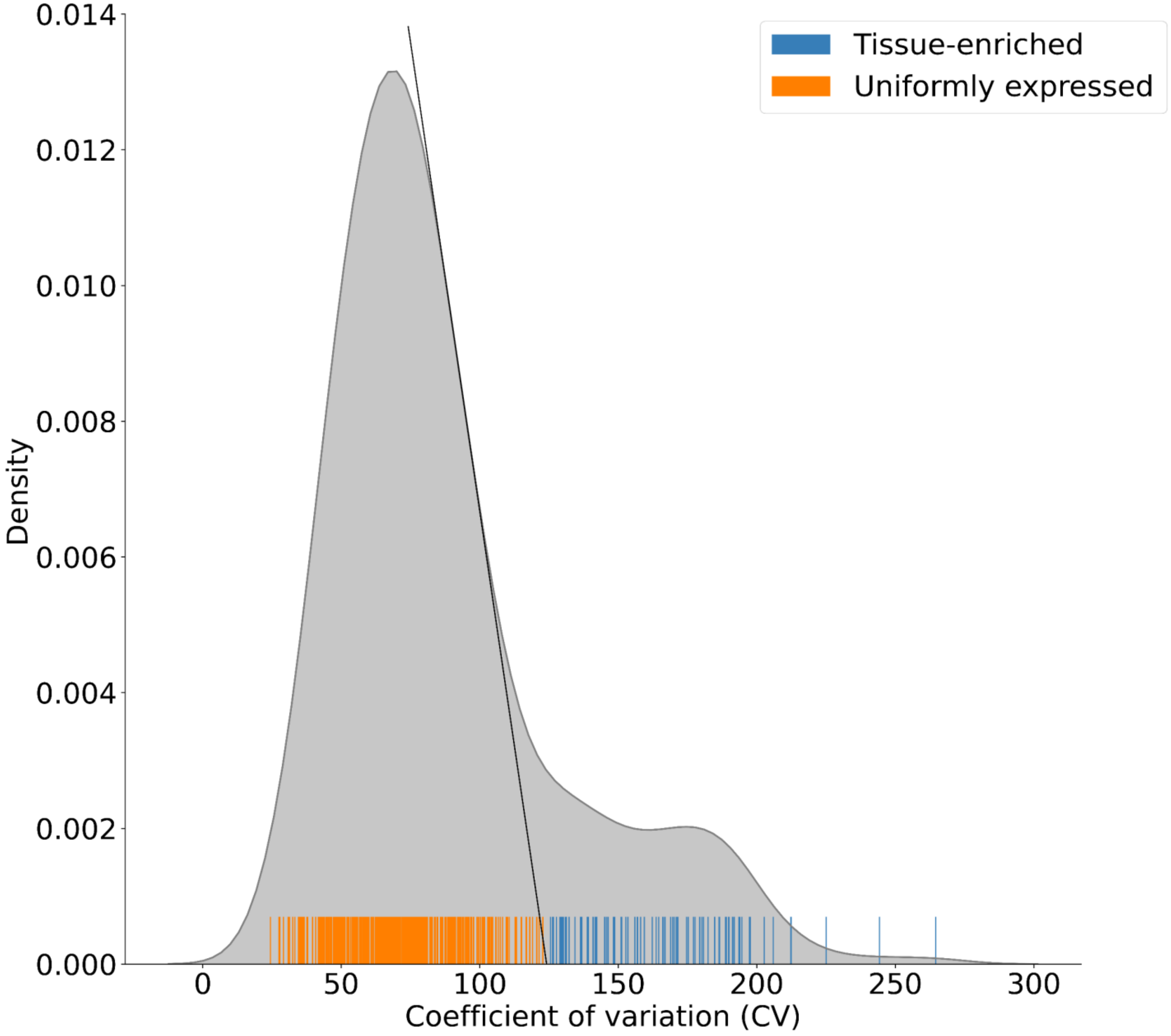


**Figure S5 (Supplementary to Figures 1 and 2)**. **Determination of the threshold used to classify the snoRNA by abundance class.** Distribution of coefficient of variation (CV) for each snoRNA between the tissues. Each line of the rugplot represented on the x-axis corresponds to the coefficient of variation of abundance of one snoRNA across all tissues. The black tangent line touches the density plot curve where its derivative is at its most negative value. The intersection of this tangent line with the x-axis represents the chosen threshold delimiting the two abundance classes which, when rounded up, corresponds to 125.

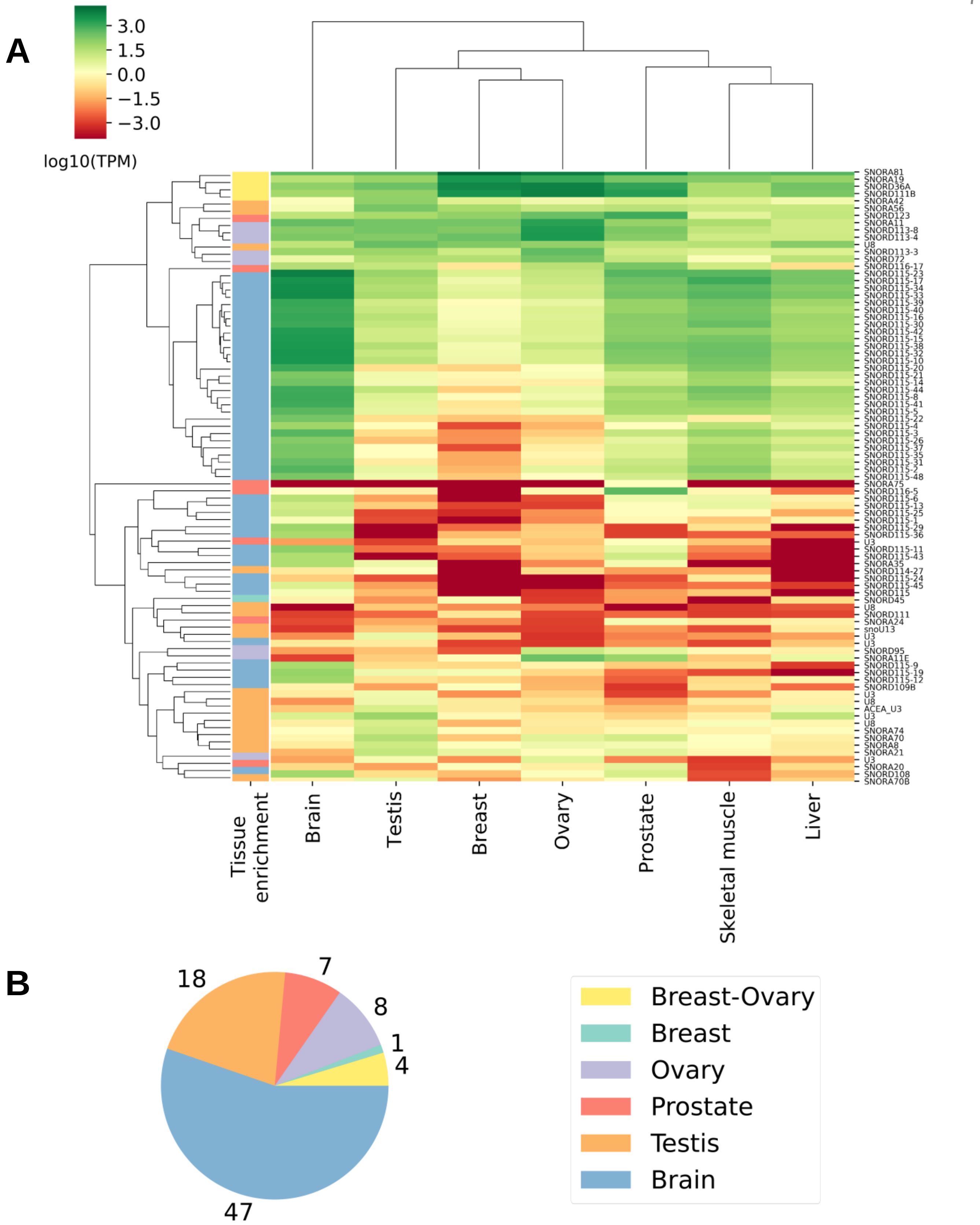


**Figure S6 (Supplementary to Figure 2). Most tissue-enriched snoRNAs are expressed in the brain.** (A) Heatmap showing the average abundance of all 85 tissue-enriched snoRNAs across the tissues (legend for tissue-enrichment color bar is shown in (B)). (B) Pie chart displaying the tissue(s) in which tissue-enriched snoRNAs are mostly expressed.

A

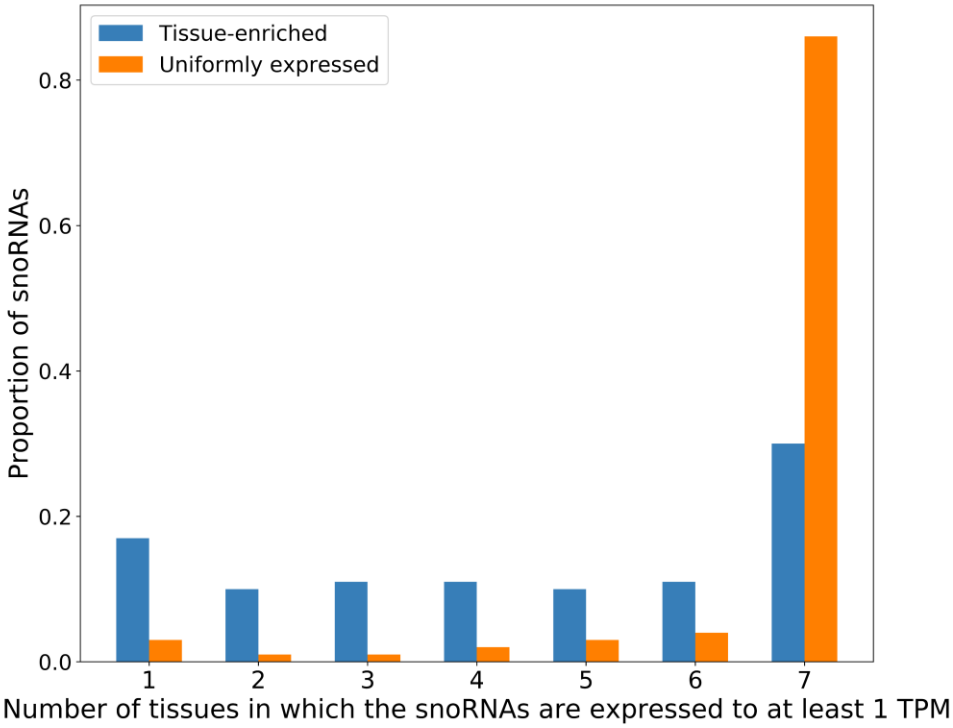


B

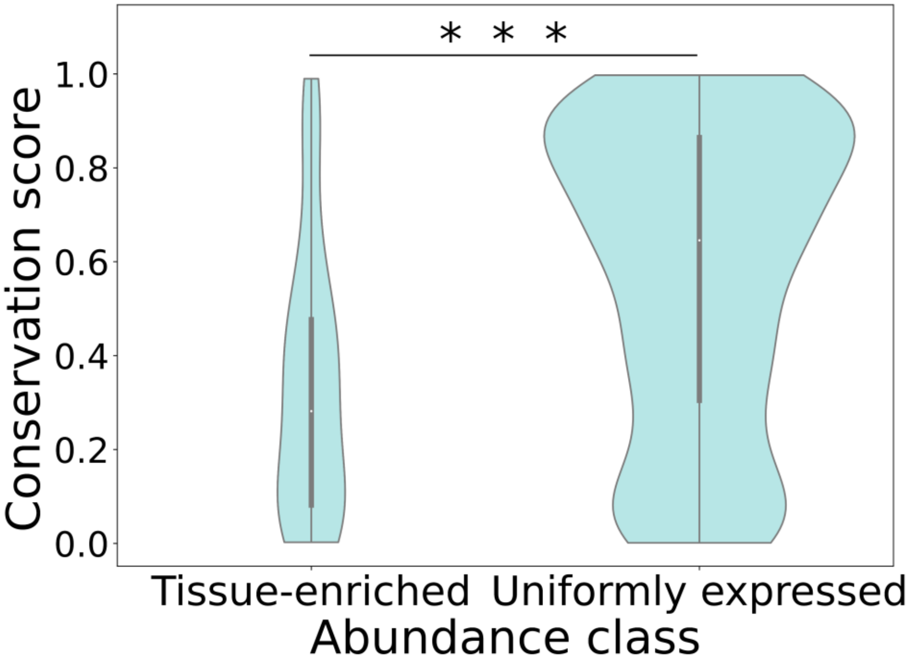


**Figure S7 (Supplementary to Figures 2 and 3). Tissue-enriched snoRNAs are less abundant in all the tissues and less conserved in primates than uniformly expressed snoRNAs.** (A) Proportion of snoRNAs classified according to their abundance class as a function of the number of tissues in which they are abundant to at least 1 TPM. (B) Violin plots showing the distribution of the phastCons conservation score in primates per abundance class. The two distributions are significantly different (Mann-Whitney U test, ***$p<4x10^{-9}$).

A

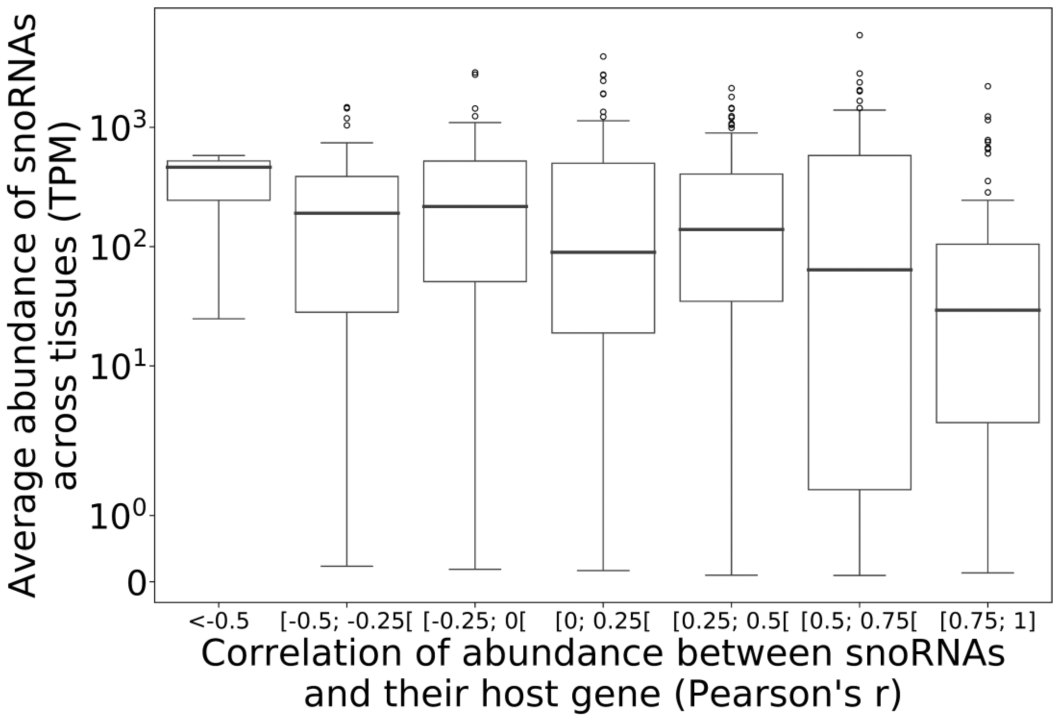


B

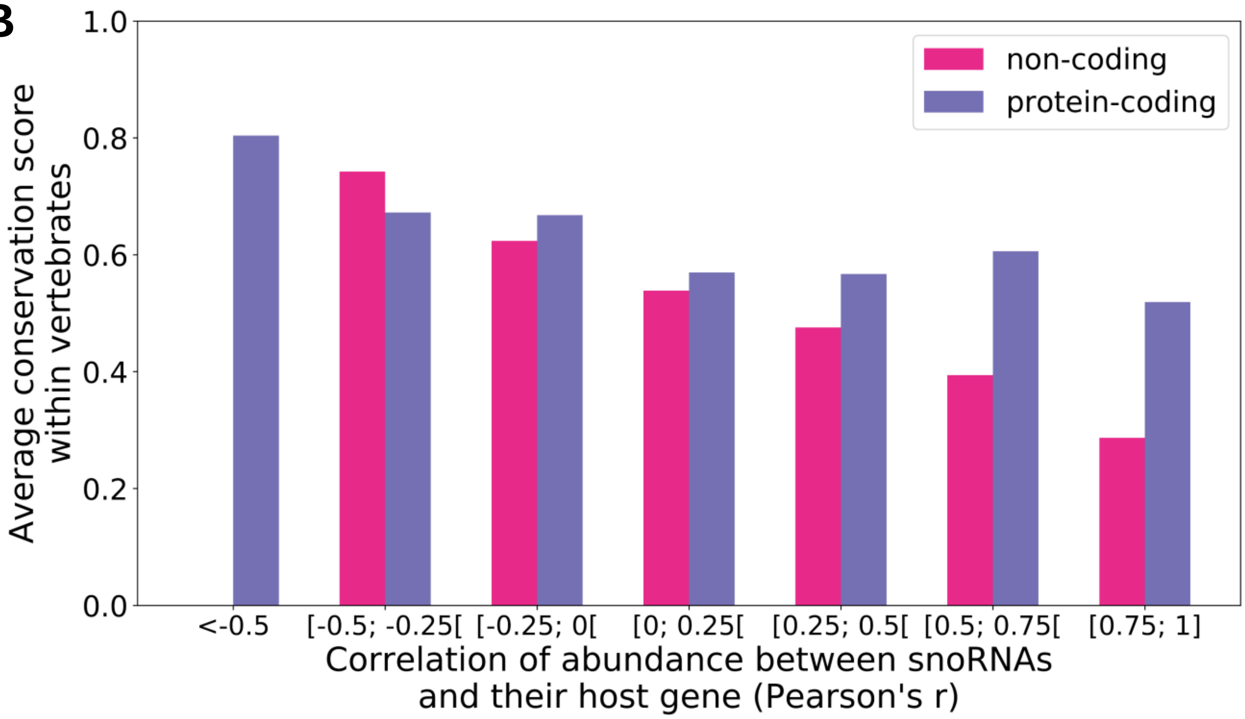


**Figure S8 (Supplementary to Figure 4)**. **Anticorrelated snoRNAs are highly abundant across tissues and conserved across vertebrates.** (A) Box plots showing the distribution of the average abundance of snoRNAs across tissues depending on the correlation of abundance with their host gene. (B) Average phastCons conservation score within vertebrates for each snoRNA as a function of the correlation of abundance with their host gene, depending on the host gene biotype.

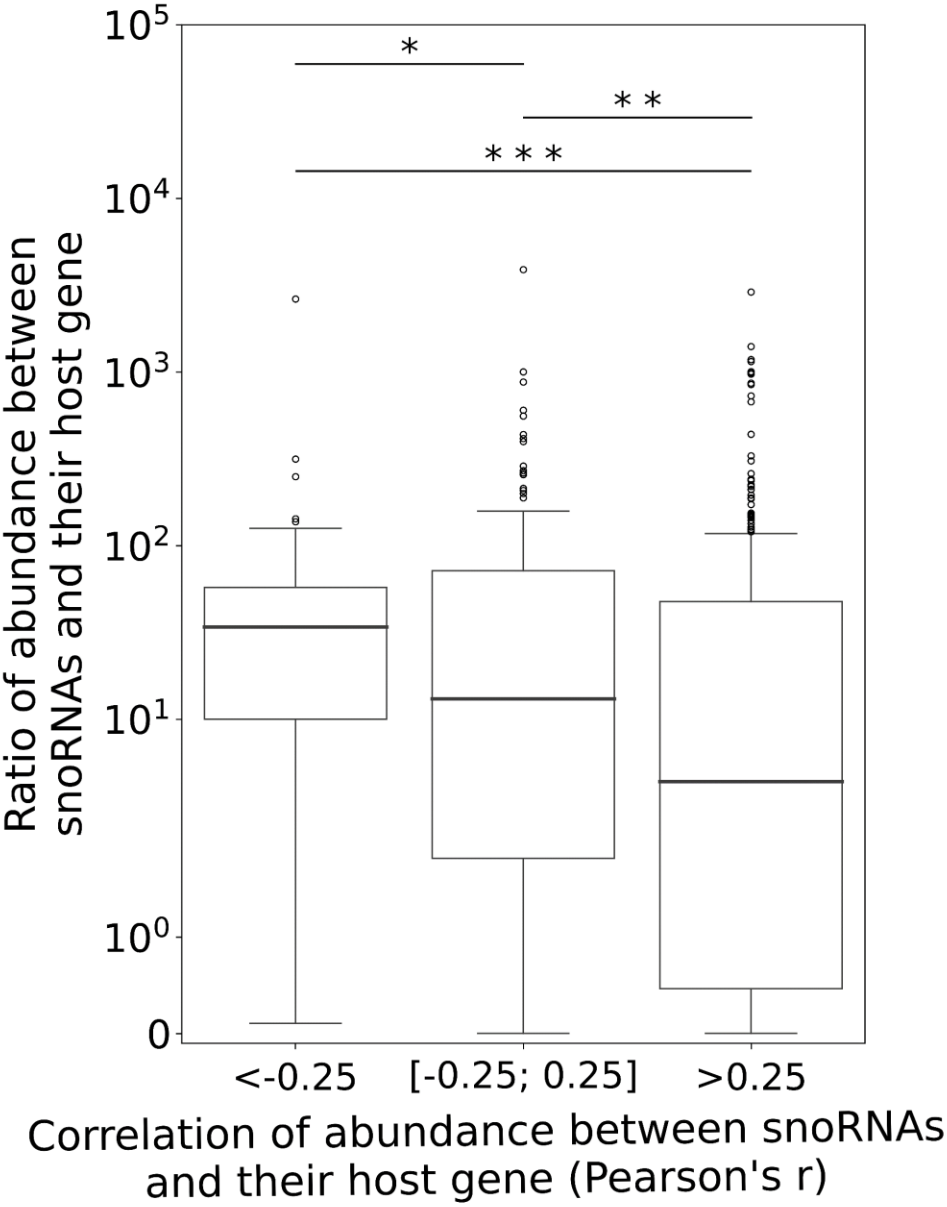


**Figure S9 (Supplementary to Figure 4). Host gene harboring anticorrelated or non-correlated snoRNAs tend to express more snoRNAs relatively to them compared to positively correlated host gene.** Box plots showing the distribution of ratios of abundance between snoRNAs and their host gene as a function of their correlation of abundance. Anticorrelated snoRNAs (Pearson's r < -0.25) are significantly more abundant relatively to their host gene compared to non-correlated (-0.25 ≤ Pearson's r ≤ 0.25) and positively correlated snoRNAs (Pearson's r > 0.25) (Mann-Whitney U test, *$p$<0.05 and ***$p$<0.0005, respectively). Non-correlated snoRNAs are also significantly more abundant relatively to their host gene compared to positively correlated snoRNAs (Mann-Whitney U test, **$p$<0.01).

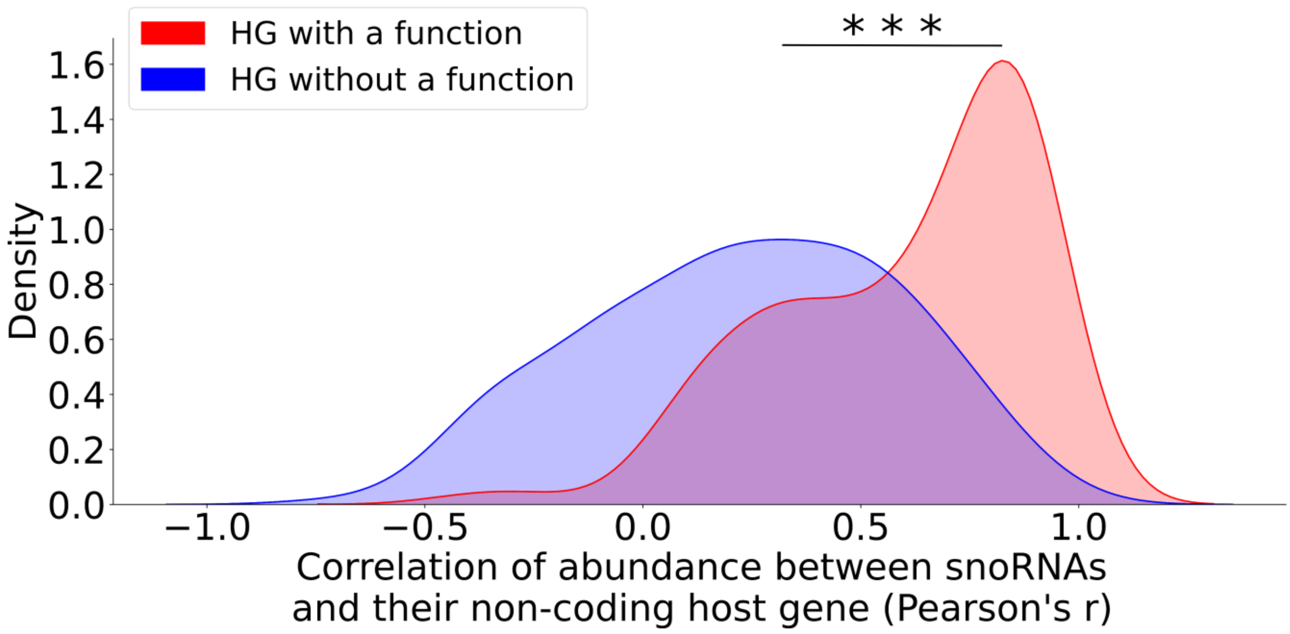


**Figure S10 (Supplementary to Figure 4). Non-coding host genes (HGs) with validated functions are more positively correlated with the expression of their snoRNAs.** Density distributions showing the correlation of abundance between snoRNAs and their non-coding HG (lncRNAs), depending on the presence or absence of snoRNA HG functionality according to lncTarD. The two distributions are significantly different (Mann-Whitney U test, ***$p$<2x$10^{-21}$).

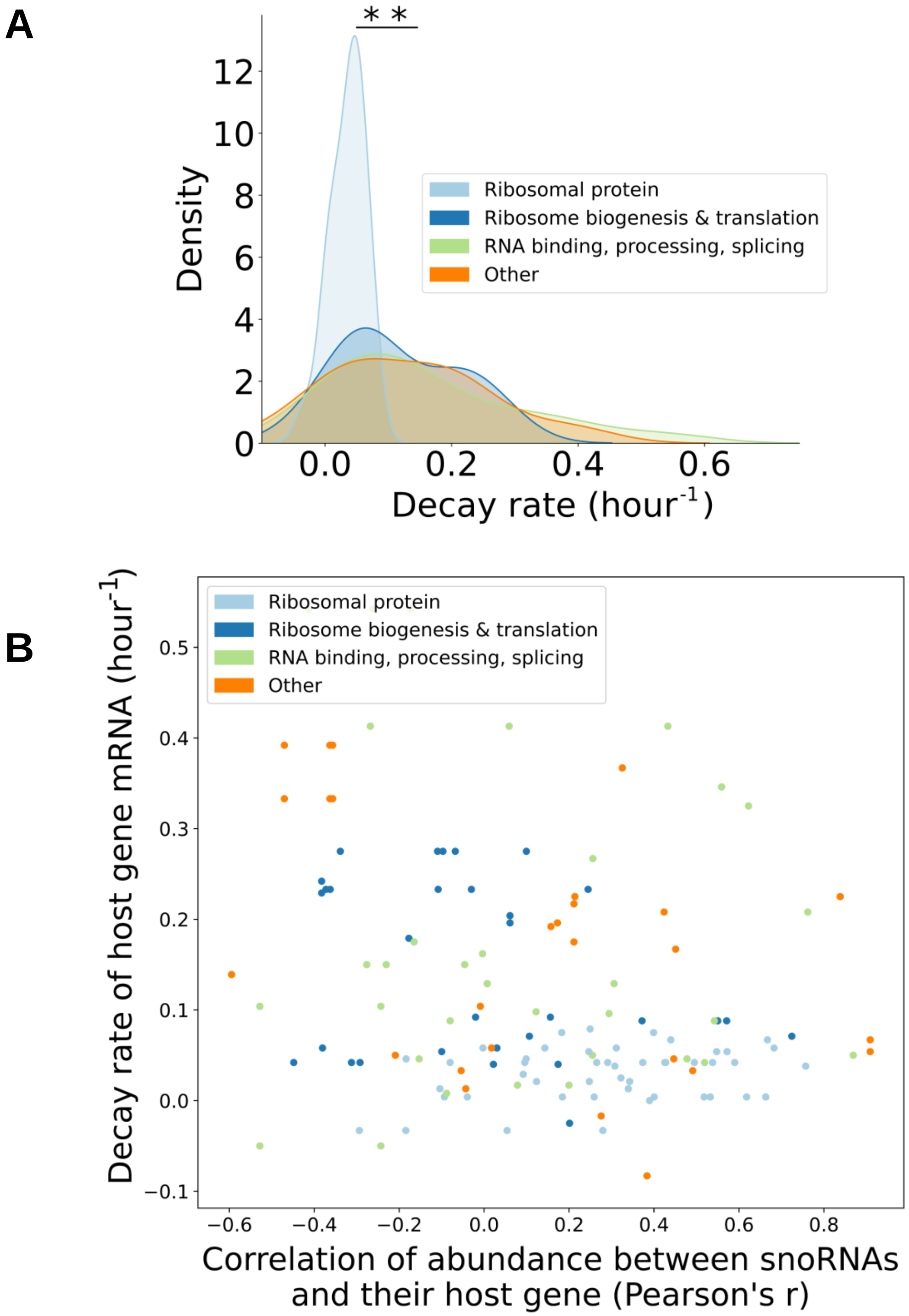


**Figure S11 (Supplementary to Figure 4). Host gene transcripts coding for ribosomal protein have lower decay rates than other types of protein-coding host gene transcripts, but the decay rate is not linked to the correlation of abundance between snoRNAs and host gene.** (A) Density distributions showing the decay rate of protein-coding host gene transcripts (decay rates retrieved from Yang et al., 2003), depending on the host gene function. The ribosomal protein distribution is significantly different than the three other distributions (Mann-Whitney U test, $^{**}p<0.01$). (B) Scatter plot displaying the decay rate of protein-coding host gene transcripts as a function of the correlation of abundance between snoRNAs and their host gene, depending on the host gene function.

**Table S1A. Top 10 most abundant protein-coding gene transcripts in the GTEx brain (frontal cortex) datasets compared to their rank of abundance in the normal brain tissue sequenced by TGIRT-Seq.**

| Gene name | GTEx rank of abundance | TGIRT-Seq rank of abundance within protein-coding RNAs in the brain† | TGIRT-Seq average rank of abundance within protein-coding RNAs in all the tissues†† | TGIRT-Seq rank of abundance within all RNAs in the brain††† |
|---|---|---|---|---|
| NRGN | 1 | 123 | 10854 | 806 |
| FTL | 2 | 32 | 86 | 585 |
| SNAP25 | 3 | 13 | 10073 | 470 |
| TMSB10 | 4 | 37 | 141 | 599 |
| GAPDH | 5 | 16 | 120 | 488 |
| CALM3 | 6 | 41 | 845 | 610 |
| ACTB | 7 | 17 | 117 | 496 |
| CALM2 | 8 | 11 | 197 | 438 |
| CKB | 9 | 66 | 3555 | 672 |
| SPARCL1 | 10 | 19 | 1193 | 507 |

**Table S1B. Top 10 most abundant protein-coding gene transcripts in the GTEx liver datasets compared to their rank of abundance in the normal liver tissue sequenced by TGIRT-Seq.**

| Gene name | GTEx rank of abundance | TGIRT-Seq rank of abundance within protein-coding RNAs in the liver† | TGIRT-Seq average rank of abundance within protein-coding RNAs in all the tissues†† | TGIRT-Seq rank of abundance within all RNAs in the liver††† |
|---|---|---|---|---|
| ALB | 1 | 1 | 7712 | 19 |
| HP | 2 | 2 | 8505 | 93 |
| SAA1 | 3 | 358 | 7646 | 1150 |
| ORM1 | 4 | 7 | 9063 | 214 |
| FTL | 5 | 21 | 86 | 304 |
| APOC3 | 6 | 11 | 11824 | 228 |
| FGB | 7 | 5 | 11555 | 191 |
| FGG | 8 | 10 | 11365 | 222 |
| APOA1 | 9 | 14 | 8151 | 238 |
| SERPINA1 | 10 | 18 | 9804 | 292 |

**Table S1C. Top 10 most abundant protein-coding gene transcripts in the GTEx breast datasets compared to their rank of abundance in the normal breast tissue sequenced by TGIRT-Seq.**

| Gene name | GTEx rank of abundance | TGIRT-Seq rank of abundance within protein-coding RNAs in the breast† | TGIRT-Seq average rank of abundance within protein-coding RNAs in all the tissues†† | TGIRT-Seq rank of abundance within all RNAs in the breast††† |
|---|---|---|---|---|
| FTL | 1 | 4 | 86 | 321 |
| FABP4 | 2 | 7 | 8587 | 401 |
| RPS18 | 3 | 17 | 36 | 477 |
| RPL13A | 4 | 26 | 61 | 537 |
| RPS12 | 5 | 22 | 45 | 516 |
| ACTB | 6 | 27 | 117 | 539 |
| RPS11 | 7 | 43 | 65 | 592 |
| EEF1A1 | 8 | 3 | 12 | 260 |
| RPL10 | 9 | 12 | 42 | 439 |
| TMSB10 | 10 | 31 | 141 | 554 |

**Table S1D. Top 10 most abundant protein-coding gene transcripts in the GTEx ovary datasets compared to their rank of abundance in the normal ovary tissue sequenced by TGIRT-Seq.**

| Gene name | GTEx rank of abundance | TGIRT-Seq rank of abundance within protein-coding RNAs in the ovary† | TGIRT-Seq average rank of abundance within protein-coding RNAs in all the tissues†† | TGIRT-Seq rank of abundance within all RNAs in the ovary††† |
|---|---|---|---|---|
| RPS18 | 1 | 8 | 36 | 509 |
| RPL13A | 2 | 17 | 61 | 572 |
| RPS12 | 3 | 16 | 45 | 562 |
| RPS11 | 4 | 24 | 65 | 623 |
| RPL10 | 5 | 5 | 42 | 490 |
| EEF1A1 | 6 | 2 | 12 | 385 |
| RPL3 | 7 | 19 | 73 | 600 |
| RPL5 | 8 | 20 | 95 | 602 |
| RPS16 | 9 | 92 | 326 | 886 |
| RPS27 | 10 | 4 | 29 | 467 |

**Table S1E. Top 10 most abundant protein-coding gene transcripts in the GTEx prostate datasets compared to their rank of abundance in the normal prostate tissue sequenced by TGIRT-Seq.**

| Gene name | GTEx rank of abundance | TGIRT-Seq rank of abundance within protein-coding RNAs in the prostate† | TGIRT-Seq average rank of abundance within protein-coding RNAs in all the tissues†† | TGIRT-Seq rank of abundance within all RNAs in the prostate††† |
|---|---|---|---|---|
| MSMB | 1 | 1 | 10879 | 95 |
| KLK3 | 2 | 2 | 11219 | 129 |
| RPL13A | 3 | 17 | 61 | 409 |
| RPS18 | 4 | 13 | 36 | 354 |
| ACTB | 5 | 36 | 117 | 502 |
| FTL | 6 | 58 | 86 | 564 |
| DES | 7 | 86 | 4660 | 652 |
| RPS11 | 8 | 27 | 65 | 468 |
| RPS12 | 9 | 22 | 45 | 443 |
| EEF1A1 | 10 | 5 | 12 | 193 |

**Table S1F. Top 10 most abundant protein-coding gene transcripts in the GTEx testis datasets compared to their rank of abundance in the normal testis tissue sequenced by TGIRT-Seq.**

| Gene name | GTEx rank of abundance | TGIRT-Seq rank of abundance within protein-coding RNAs in the testis† | TGIRT-Seq average rank of abundance within protein-coding RNAs in all the tissues†† | TGIRT-Seq rank of abundance within all RNAs in the testis††† |
|---|---|---|---|---|
| PRM2 | 1 | 5 | 11478 | 173 |
| PRM1 | 2 | 2 | 11594 | 146 |
| TNP1 | 3 | 6 | 11667 | 198 |
| PTGDS | 4 | 29 | 4149 | 540 |
| PHF7 | 5 | 9 | 8703 | 359 |
| LELP1 | 6 | 390 | 12597 | 1242 |
| TUBB4B | 7 | 30 | 1114 | 541 |
| GPX4 | 8 | 21 | 322 | 494 |
| SMCP | 9 | 101 | 12439 | 751 |
| CMTM2 | 10 | 42 | 11518 | 583 |

**Table S1G. Top 10 most abundant protein-coding gene transcripts in the GTEx skeletal muscle datasets compared to their rank of abundance in the normal skeletal muscle tissue sequenced by TGIRT-Seq.**

| Gene name | GTEx rank of abundance | TGIRT-Seq rank of abundance within protein-coding RNAs in the skeletal muscle† | TGIRT-Seq average rank of abundance within protein-coding RNAs in all the tissues†† | TGIRT-Seq rank of abundance within all RNAs in the skeletal muscle††† |
|---|---|---|---|---|
| ACTA1 | 1 | 20 | 10925 | 285 |
| CKM | 2 | 2 | 10510 | 74 |
| MYL2 | 3 | 16 | 11901 | 258 |
| DES | 4 | 5 | 4660 | 106 |
| MB | 5 | 6 | 10121 | 161 |
| TNNC2 | 6 | 9 | 10077 | 185 |
| TNNC1 | 7 | 10 | 10357 | 204 |
| GAPDH | 8 | 11 | 120 | 205 |
| TNNT1 | 9 | 31 | 10273 | 328 |
| TPM2 | 10 | 21 | 2616 | 287 |

† This rank corresponds to the corresponding rank for the top 10 protein-coding genes identified by GTEx within the protein-coding RNAs detected by TGIRT-Seq in the specified tissue. There is an average difference of only 27 ranks out of a total of 14 914 protein-coding RNAs considered when comparing this rank with the GTEx ranking, confirming the accuracy of the TGIRT-Seq quantification not only for non-coding RNAs, but also for protein-coding transcripts.

†† This rank corresponds to the average rank across all the tissues used in the TGIRT-Seq (not only in the specified tissue) and within protein-coding RNAs. This rank is on average 4 499 ranks lower (i.e. farther away from the most abundant transcripts) than the average TGIRT-Seq rank in the specified tissue and within protein-coding RNAs (comparison between the fourth and third columns respectively), confirming the validity and identity of the tissues we used in the TGIRT-Seq.

††† This rank corresponds to the corresponding rank for the top 10 protein-coding genes identified by GTEx within all RNAs detected by TGIRT-Seq in the specified tissue (not only the protein-coding RNAs). This rank is on average 341 ranks lower (i.e. farther away from the most abundant transcripts), supporting the notion of the human transcriptome being dominated by specific subsets of highly abundant non-coding RNAs (Boivin et al., 2018).

**Table S2. Proportion of expressed RNA in the TGIRT-Seq datasets.**

| RNA biotype | Number of expressed RNA (>1 TPM in at least one sample) | Total number of annotated RNA | Proportion of expressed RNA in the TGIRT-Seq datasets (%) |
|---|---|---|---|
| snoRNA | 475 | 947 | 50 |
| snRNA | 451 | 1900 | 24 |
| tRNA | 545 | 651 | 84 |
| Protein-coding RNA | 14927 | 20367 | 73 |
| lncRNA | 2493 | 14703 | 17 |

**Table S3. Sequencing depth and quality assessment of the TGIRT-Seq datasets generated for each tissue.**

| | Breast average | Ovary average | Prostate average | Testis average | Liver average | Skeletal muscle average | Brain average |
|---|---|---|---|---|---|---|---|
| Total number of reads | 62 086 000 | 58 077 456 | 68 666 341 | 60 247 563 | 58 656 576 | 77 793 106 | 57 189 594 |
| Average read length (nt) | 133 | 126 | 127 | 121 | 124 | 125 | 126 |
| Uniquely mapped reads (%) | 77,7 | 71,9 | 78,6 | 79,3 | 74,2 | 66,3 | 76,6 |
| Multi-mapped reads (%) | 21,4 | 27,4 | 20,6 | 19,9 | 25,1 | 32,0 | 22,7 |
| Unmapped reads (%) | 0,9 | 0,7 | 0,8 | 0,8 | 0,7 | 1,7 | 0,6 |

**Table S4. Donor information and sample quality for each tissue sample.**

| Sample name | Sex | Age | RNA integrity number (RIN) |
|---|---|---|---|
| Breast_1 | Female | N/A | 8.4 |
| Breast_2 | Female | N/A | 9.4 |
| Breast_3 | Female | N/A | 8.4 |
| Ovary_1 | Female | N/A | 7.1 |
| Ovary_2 | Female | N/A | 8.1 |
| Ovary_3 | Female | N/A | 7.0 |
| Prostate_1 | Male | 68 | 9.0 |
| Prostate_2 | Male | 63 | 8.9 |
| Prostate_3 | Male | 71 | 9.1 |
| Testis_1 | Male | 44 | 7.9 |
| Testis_2 | Male | 29 | 7.1 |
| Testis_3 | Male | 64 | 8.0 |
| Liver_1 | Female | 30 | 5.4 |
| Liver_2 | Male | 75 | 7.2 |
| Liver_3 | Male | 29 | 8.2 |
| Brain_1 | Male | 41 | 7.1 |
| Brain_2 | Male | 24 | 7.9 |
| Brain_3 | Male | 27 | 7.9 |
| Skeletal_muscle_1 | Male | 43 | 9.2 |
| Skeletal_muscle_2 | Male | 76 | 9.4 |
| Skeletal_muscle_3 | Male | 24 | 9.3 |
